# Spatial control of genome editing activity enables localized immunotherapy

**DOI:** 10.64898/2026.01.14.699490

**Authors:** Xiaoyue Yang, Laura Tong, Yidan Pan, Jin Huang, Zhongchao Yi, Daheng He, Jingpeng Liu, Chi Wang, Ying Liang, Sheng Tong

## Abstract

Precise control of genome editing activity in vivo remains a major barrier to the clinical translation of CRISPR-based therapeutics, as current approaches cannot reliably restrict editing activity to target tissues ^1,2,3^. Here we show that a magnetically gated baculoviral CRISPR platform enables spatial control of genome editing activity ^4^. By coupling magnetic activation of viral transduction at target sites with complement-mediated inactivation of circulating vectors, this system functionally confines genome editing activity to defined spatial contexts, effectively decoupling editing activity from biodistribution. We further show that spatially confined genome editing can be coupled with endogenous antiviral sensing to drive coordinated immune modulation within tumours. In a syngeneic murine tumour model, local administration combined with magnetic activation restricts genome editing to tumours without detectable editing in major organs, remodels dendritic cell and CD8□ T cell states, suppresses tumour growth, and extends survival. Together, these findings establish a generalizable strategy for spatial control of genome editing and localized immunomodulation in cancer.

---

Achieving such spatial control of genome editing activity in vivo requires overcoming two fundamental challenges: the inability to restrict vector transduction to target tissues and the lack of mechanisms to suppress editing activity following systemic dissemination. Existing delivery systems, including viral vectors and lipid nanoparticles, can improve targeting but remain systemically distributed following both local and systemic administrations, with vectors frequently detected beyond the site of delivery ^5,6,7^. This leads to off-target accumulation and sustained nuclease activity, increasing the risk of unintended genome editing and genotoxicity-associated adverse effects ^8,9,10^. As a result, editing outcomes are primarily dictated by biodistribution rather than by controllable activation, thereby limiting the safe and precise use of CRISPR-based therapeutics in vivo.

Baculovirus provides a programmable viral delivery backbone whose biological properties can be leveraged to impose spatial control over genome editing activity. Although replication-defective in mammalian cells, baculoviral vectors support large genetic payloads and efficient nuclear delivery, enabling multiplexed CRISPR-mediated genome editing ^11,12,13^. Importantly, following in vivo administration, baculoviral particles are rapidly inactivated by complement, restricting systemic persistence but also limiting their utility as gene therapy vectors ^14^. We previously established magnetically controlled and complement-constrained baculoviral genome editing in vivo ^4^. In this framework, magnetic fields enhance local viral entry, whereas complement activity suppresses transduction outside the activated region, jointly defining where genome editing can occur. Outside the magnetic field, complement-mediated inactivation restricts vector activity, establishing orthogonal physical and biological constraints in vivo. However, whether this control architecture can be translated into therapeutically effective genome editing remains unclear. Here we develop a magnetically gated baculovirus (MBV) platform for localized cancer immunotherapy that integrates spatially gated genome editing with endogenous immune regulation, extending prior magnetic control strategies (Fig. 1).

Immune checkpoint blockade can elicit durable antitumour responses, yet tumour heterogeneity and adaptive resistance often necessitate combination strategies ^15,16,17,18^. These approaches remain constrained by incomplete coordination of immunomodulation within tumours and systemic immune perturbation ^19^. Strategies that enable localized control of immune regulatory feedback within tumours could expand the therapeutic window of combination immunotherapy. Oncolytic viruses leverage antiviral immune activation to stimulate antitumour immunity; however, viral sensing induces interferon-driven programs that both promote immune activation and upregulate PD-L1, creating a compensatory feedback loop that restrains effector function ^20,21,22^. Although genome editing could, in principle, resolve this inhibitory feedback locally, incorporating gene disruption into replicating viral platforms introduces safety concerns related to uncontrolled dissemination and persistent nuclease activity.

Likewise, unmethylated CpG motifs within the baculoviral genome engage innate immune sensors and induce interferon-driven programs that reshape local immune environments ^23,24^. We therefore sought to harness this intrinsic antiviral response while resolving its associated inhibitory feedback. We engineer a genome editing platform that resolves antiviral-induced checkpoint feedback locally by coupling viral delivery with spatially gated CRISPR-mediated gene disruption. Genome editing provides a programmable route to durable tumour immunomodulation, whereas magnetic fields provide a non-invasive means to spatially control therapeutic activation. Notably, clinical studies of magnetically guided nanoparticle targeting in tumours have demonstrated that externally applied magnetic fields can localize therapeutic agents within deep tissues ^25,26^, supporting the translational feasibility of magnetically gated therapeutic strategies. This integrated architecture enables spatially restricted genome editing while preserving virus-induced innate immune amplification.

This approach establishes control of genome editing activity—not vector delivery—as the primary determinant of therapeutic specificity in vivo. To evaluate whether spatially gated genome editing can resolve tumour-intrinsic immune feedback in vivo, we applied this platform to disrupt *Pdl1* within solid tumours. In a syngeneic colon cancer model with modest responsiveness to PD-L1 antibody blockade, magnetic activation enabled tumour-restricted CRISPR-mediated checkpoint disruption without detectable editing in normal tissues. Tumour-restricted editing resulted in tumour growth suppression and extended survival. Together, these results establish a genome-editing architecture that integrates physical gating, biological containment, and programmable checkpoint disruption to enable localized combination immunotherapy.

## Magnetic activation restores baculoviral transduction under complement-mediated constraint

We first examined whether magnetic activation could overcome complement-mediated restriction of baculoviral transduction in MC38 murine colon adenocarcinoma cells, the cell line used to establish the syngeneic tumour model. MBV was generated by electrostatic conjugation of MNPs to the surface of baculoviral vectors (BVs). MNPs functionalized with TAT peptides were synthesized using a two-step procedure that enabled control over particle size, dispersity, and peptide density (Fig. 2a, Supplementary Fig. S1) ^27^. The cationic TAT peptides facilitated binding to the negatively charged baculoviral envelope. BV vectors encoding SpCas9, reporters, and gRNAs were constructed within a single viral genome, leveraging the large cargo capacity of BV to improve editing efficiency and reduce production complexity (Supporting Fig. S2). MBV complexes were assembled by mixing MNPs with BV at a ratio of ∼3 µg Fe per 10^7^ pfu for 10 min in PBS (Fig. 2a).

**Fig. 1.**
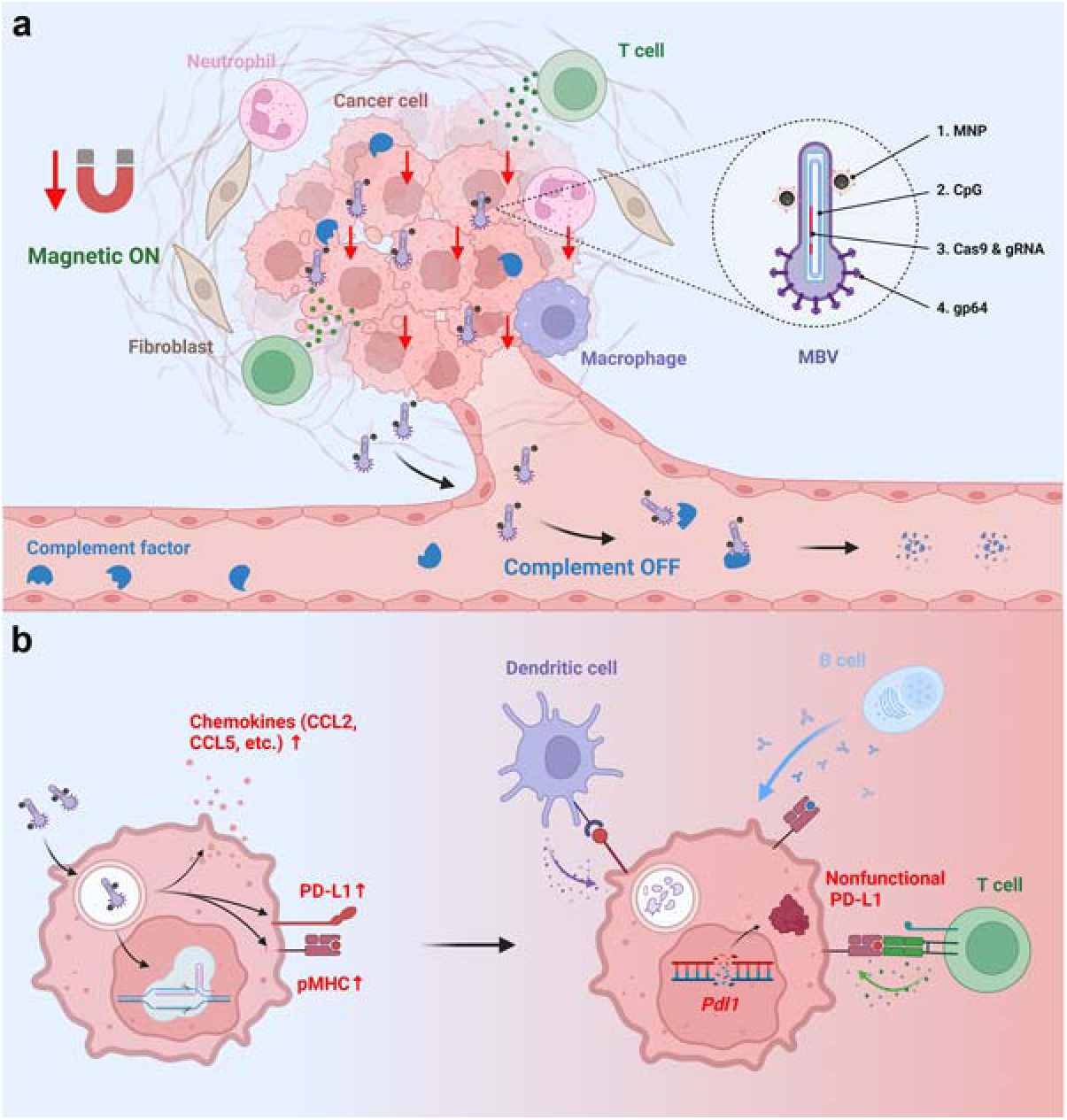
Magnetically gated control of genome editing activity enables localized immunotherapy. **a**. MBVs are administered by intratumoural infusion followed by brief magnetic activation. Magnetic activation functions as a physical on-switch that enhances local viral entry and genome editing activity at target sites despite complement activity, thereby confining genome editing activity to magnetically targeted tumour regions. MBVs that enter the systemic circulation are rapidly inactivated by complement, which functions as a biological off-switch to suppress genome editing activity outside the activated region, limiting dissemination and off-target exposure. Inset: Structural components of MBV include: (1) magnetic nanoparticles (MNPs), which enable magnetic responsiveness; (2) intrinsic CpG motifs within the baculoviral genome, which trigger antiviral sensing; (3) Cas9 and guide RNA expression cassettes, which enable target gene disruption; and (4) gp64, which mediates viral entry and is sensitive to complement inactivation. **b**. MBV transduction activates innate immune sensing pathways, inducing chemokine secretion and enhancing antigen presentation within tumour cells while concurrently driving compensatory PD-L1 upregulation. CRISPR-mediated disruption of *Pdl1* eliminates PD-L1-mediated inhibitory feedback, sustaining effector cell activation in the tumour microenvironment. Together, spatially confined genome editing activity and antiviral immune activation integrate to drive localized antitumour immunity.

**Fig. 2.**
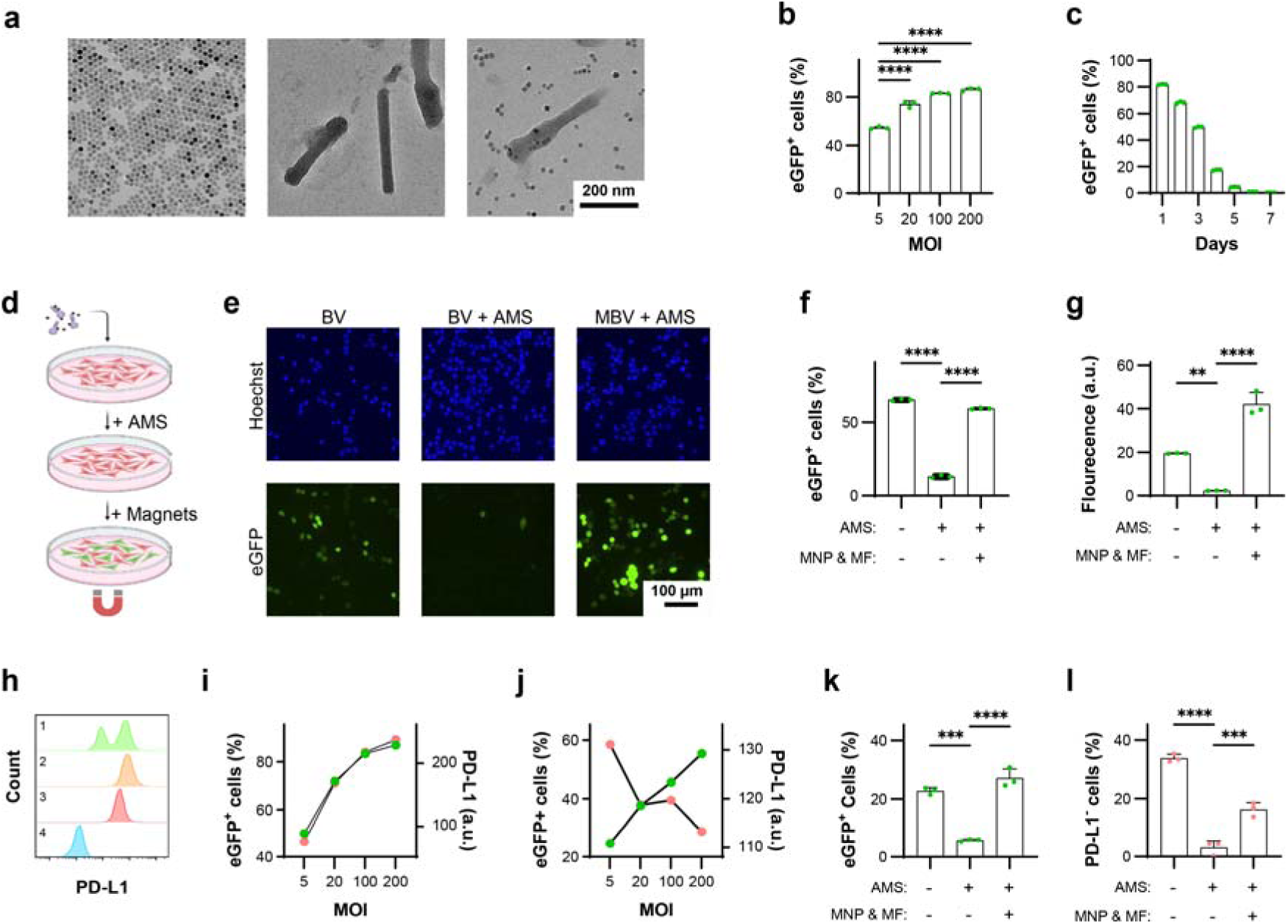
Characterization of MBV. **a**. Transmission electron microscopy images of magnetic nanoparticles (MNPs), baculovirus (BV), and magnetically modified baculovirus (MBV). Samples in the two right panels were negatively stained with phosphotungstic acid. **b**, **c**. Efficiency and temporal profile of BV transduction in MC38 cells. For panel **c**, cells were transduced at a multiplicity of infection (MOI) of 20. **d.** Schematic of the in vitro assay used to evaluate magnetic activation of MBV–eGFP under complement pressure. **e**. Representative fluorescence microscopy images of MC38 cells incubated with BV–eGFP alone, BV–eGFP in the presence of adult mouse serum (AMS), and MBV–eGFP with AMS and magnetic activation. **f**, **g**. Flow cytometry analysis of AMS-mediated inhibition and magnetic activation of BV–eGFP-driven eGFP expression. **h**. Representative flow cytometry histograms of PD-L1 expression in (**1**) MC38 cells transduced with BV–*Pdl1*–eGFP (MOI = 20), (**2**) MC38 cells transduced with BV–eGFP (MOI = 20), (**3**) wild-type MC38 cells, and (**4**) isotype control-stained MC38 cells. **i**, **j**. Quantification of eGFP (green) and PD-L1 (pink) expression levels in MC38 cells transduced with BV–eGFP and BV–*Pdl1*–eGFP, respectively. **k**, **l**. Flow cytometry analysis of AMS inhibition and magnetic activation of BV–*Pdl1*–eGFP-mediated eGFP expression and PD-L1 disruption in MC38 cells. Data are presented as mean ± s.d. ns, ***, and **** denote not significant, *P* < 0.001, and *P* < 0.0001, respectively.

BV transduction efficiency was first evaluated in MC38 cells using BV–eGFP. More than 50% of cells expressed eGFP at a multiplicity of infection (MOI) of 5 (Fig. 2b). eGFP expression peaked at 24 h and declined to undetectable levels by day 7 (Fig. 2c), consistent with transient BV-mediated transgene expression. To assess complement inhibition and magnetic activation as functional off- and on-switches in MC38 cells, BV transduction was evaluated in the presence of 50% adult mouse serum (AMS), which contains a fully developed complement system (Fig. 2d, Supplementary Fig. S3). AMS markedly reduced both the fraction of eGFP-positive cells and expression intensity (Fig. 2e). In contrast, magnetic activation of MBV restored robust transduction under complement pressure, yielding higher expression levels than BV in AMS-free conditions. Flow cytometry showed ∼80% inhibition of BV transduction by AMS, whereas magnetically activated MBV fully rescued transgene expression (Fig. 2f, g). These results confirm that complement sensitivity suppresses BV activity, while magnetic activation restores BV-mediated gene delivery.

The murine *Pdl1* locus comprises five exons on chromosome 19. Six candidate guide RNAs targeting early exons or key structural regions were designed and screened in MC38 cells (Supplementary Table S1) ^28^. Based on T7E1 cleavage and next-generation sequencing (NGS) analysis, a guide targeting exon 3 was selected (Supplementary Fig. S4). NGS revealed that ∼87% of Cas9-induced indels were frameshift mutations, indicating efficient functional disruption.

Consistent with observations in oncolytic viruses, BV–eGFP transduction increased PD-L1 expression in MC38 cells in a transduction-dependent manner, reflecting interferon-mediated checkpoint induction (Fig. 2h, i) ^20^. In contrast, BV–*Pdl1*–eGFP reduced PD-L1 expression at matched MOIs without inducing substantial cytotoxicity (Fig. 2j, Supplementary Fig. S5). MBV–*Pdl1* achieved greater disruption efficiency than BV–*Pdl1* (Supplementary Fig. S6). AMS suppressed both eGFP expression and *Pdl1* disruption by BV–*Pdl1*–eGFP, whereas magnetic activation of MBV–*Pdl1*–eGFP partially restored editing activity under complement pressure (Fig. 2k, l). Together, these findings establish complement inactivation and magnetic activation as orthogonal biological and physical constraints that regulate vector activity in vivo.

## BV transduction induces immune-associated signalling in vitro

To isolate immune effects attributable to BV transduction independent of *Pdl1* disruption, a control vector encoding Cas9 without a targeting guide RNA (BV–Cas9) was generated. Similar to BV–eGFP, BV–Cas9 increased PD-L1 expression relative to untreated cells (Supplementary Fig. S7).

Transcriptomic profiling of MC38 cells treated with BV–*Pdl1*, MBV–Cas9, and MBV–*Pdl1* was performed using the nCounter® Tumor Signaling 360 Panel. Both BV–*Pdl1* and MBV–*Pdl1* reduced *Pdl1* transcript level (Fig. 3a). In contrast, global transcriptional changes were largely shared across all treated groups and distinct from untreated controls, indicating that BV transduction was the dominant driver of immune-related gene expression (Fig. 3b-d). BV-treated cells exhibited marked upregulation of chemokines associated with lymphocyte recruitment, including *Ccl2*, *Cxcl11*, *Cxcl10*, and *Ccl5* (Fig. 3e, Supplementary Fig. S8a, c). Gene-set analysis revealed activation of chemokine signalling, antigen presentation, and interferon response pathways in MBV–Cas9 and MBV–*Pdl1* but to a lesser extent in the BV–*Pdl1* group (Fig. 3f, Supplementary Fig. S8b, d), consistent with a coordinated antiviral response enhanced magnetically. All BV-treated cells also exhibited activation of T-cell costimulation pathways.

**Fig. 3.**
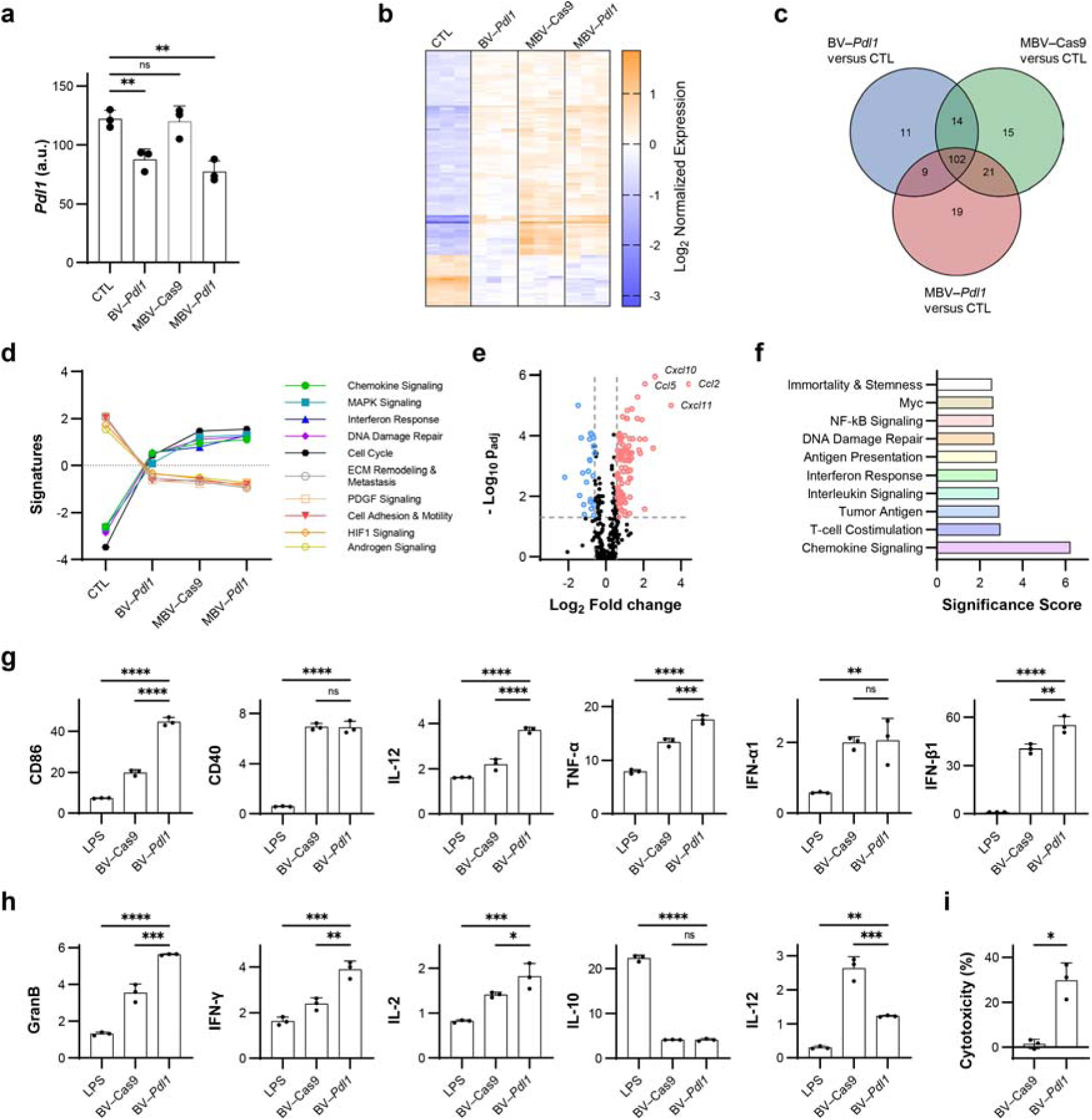
BV–*Pdl1* enhances anticancer immunity through coordinated antiviral responses and *Pdl1* disruption. Transcriptomic responses of MC38 cells treated with BV–*Pdl1*, MBV–Cas9, and MBV–*Pdl1* were analyzed using the nCounter® Tumor Signaling 360 Panel. **a**. *Pdl1* expression in MC38 cells following the indicated treatments. ns and ** denote no significance and *P* < 0.01, respectively. **b**. Heat map of normalized gene expression across treatment groups. n = 3 per group. **c**. Venn diagram of differentially expressed genes (|log_2_FC| > 0.58, *P*_adj_ < 0.05) in BV–*Pdl1*-, MBV–Cas9-, MBV–*Pdl1*-treated cells relative to untreated controls. **d.** Pathway analysis. The pathway scores are fit using the first principal component of each gene set’s data. Lines show each pathway’s average score across values of each group (n = 3). **e**. Differential gene expression in MC38 cells treated with MBV–*Pdl1* relative to PBS control. *P* values were adjusted using the Benjamini and Yekutieli method for multiple testing correction. **f**. Pathway enrichment analysis of MC38 cells treated with MBV–*Pdl1* relative to PBS control. Bone marrow-derived dendritic cells (BMDCs) were cocultured with BV–*Pdl1*-treated MC38 cells. **g**. Gene expression in BMDCs quantified by RT-qPCR and normalized to coculture with untreated MC38 cells. Lipopolysaccharide (LPS) treatment was used as a positive control (n = 3 per group). Splenocytes were cocultured with BV–*Pdl1*-treated MC38 cells. **h**. Gene expression in splenocytes quantified by RT-qPCR and normalized to coculture with untreated MC38 cells. LPS treatment was used as a positive control (n = 3 per group). **i**. Splenocytes-mediated cytotoxicity against BV-transduced MC38 cells relative to untreated control (n = 3 per group). Data are presented as mean ± s.d. ns, *, **, ***, and **** denote no significance, *P* < 0.05, *P* < 0.01, *P* < 0.001, and *P* < 0.0001, respectively.

To evaluate downstream immune activation, BV-transduced MC38 cells were cocultured with bone marrow-derived dendritic cells (BMDCs). Coculture induced upregulation of canonical maturation markers (*Cd86*, *Cd40*, *Il-12*, *Tnf-α*, *Ifn-α1*, and *Ifn-β1*), indicating dendritic cell activation (Fig. 3g). When cocultured with splenocytes, splenocytes exposed to BV–*Pdl1*-treated cells exhibited elevated expression of cytotoxic effector genes, including *Gzmb*, *Ifn-γ*, relative to BV–Cas9 (Fig. 3h). Furthermore, splenocytes exhibited significant toxicity to BV–*Pdl1*-treated MC38 cells but not BV–Cas9-treated cells (Fig. 3i). These findings indicate that disruption of *Pdl1* mitigates antiviral-induced compensatory PD-L1 upregulation, coupling immune recruitment with checkpoint inhibition to sustain cytotoxic effector activity.

## Magnetic gating enables functional confinement of genome editing in vivo

To evaluate whether localized *Pdl1* disruption could modulate tumour immunity in vivo, C57BL/6 mice bearing established MC38 tumours (∼50 mm^3^) were treated by intratumoural infusion of MBV–*Pdl1* followed by magnetic activation. This experimental design enabled spatially confined genome editing within established tumours under conditions that model intervention in clinically accessible lesions.

Magnetic force on MBV is governed by field gradients rather than field strength alone (see Methods). Prior studies have demonstrated that high field gradients can be generated at substantial depth within the mouse body ^29,30^. To achieve reproducible magnetic activation while intentionally limiting force exposure in major organs, we designed a magnet array composed of four cylindrical NdFeB magnets arranged in alternating polarity and centered around the tumour (Fig. 4a). The magnet array was integrated into a 3D-printed animal bed to ensure consistent tumour positioning (Supplementary Fig. S9). Following vector infusion, mice were maintained under anesthesia with tumours aligned within the magnetic field for 1 h. Simulated magnetic flux density and force profiles at the tumour surface were comparable to those used for in vitro activation (Fig. 4b, c, Supplementary Fig. S10), confirming that the in vivo configuration provided sufficient magnetic gating to trigger MBV activation. Estimated forces on MNPs in the liver and spleen were approximately two orders of magnitude lower than those at the tumour site. This confined magnetic profile reflects intentional array design to localize activation, rather than a limitation in field penetration.

**Fig. 4.**
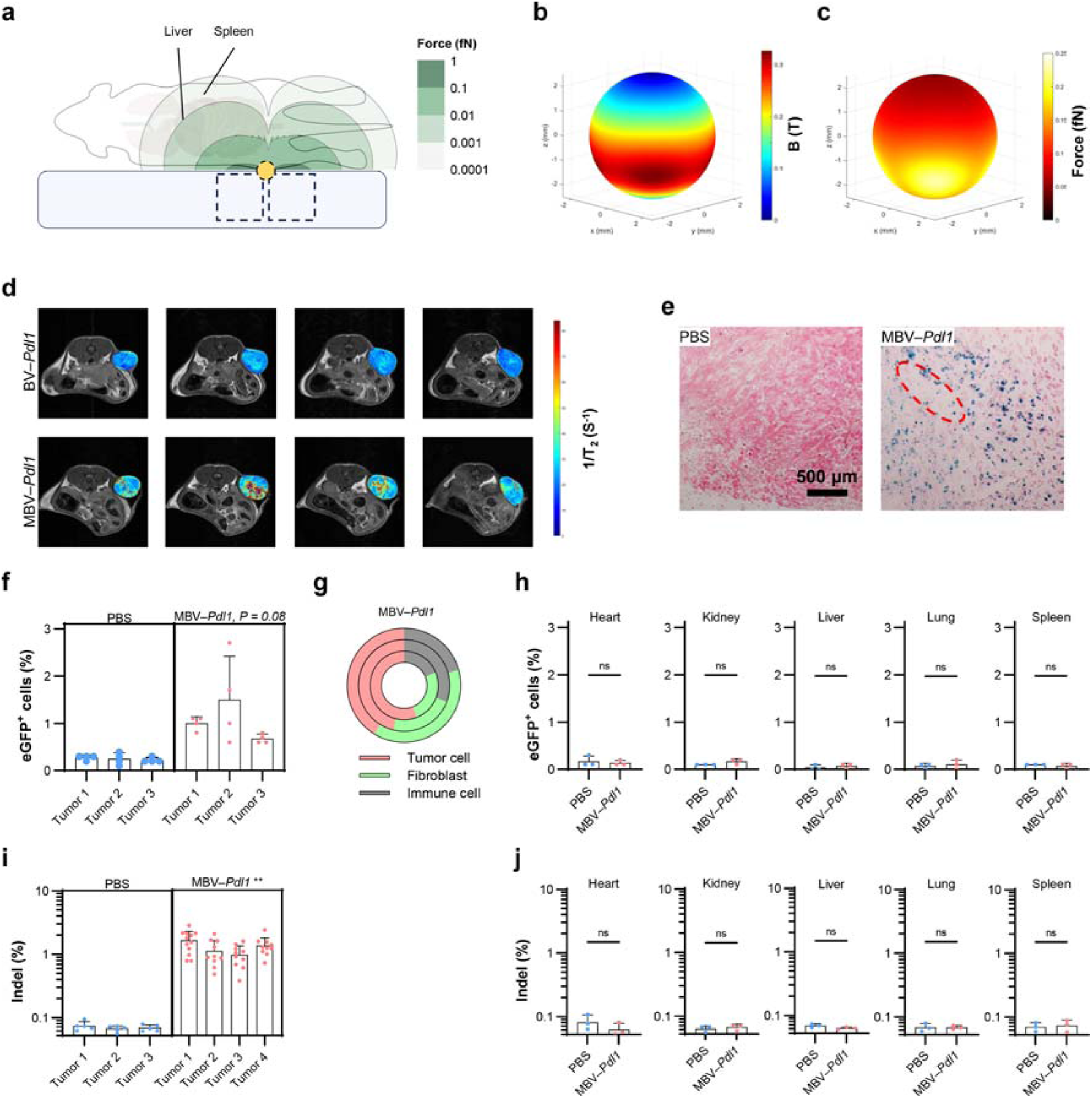
Magnetic gating enables functional confinement of genome editing in vivo. **a**. Schematic of in vivo magnetic activation. A mouse outline is overlaid with simulated magnetic force contours generated by the cylindrical magnet array (dashed boxes). A subcutaneous tumour (yellow) resides within the high-force region. Contour levels (1 to 10□□ fN) demonstrate the rapid spatial decay of magnetic force, supporting localized activation at the tumour site while limiting exposure to major organs (liver and spleen). The magnet configuration is designed to confine magnetic force to the tumour region rather than reflecting intrinsic limits of field penetration. The schematic is drawn to scale. **b**, **c**. Simulated distributions of magnetic flux density (**b**) and magnetic force (**c**) at the tumour surface; force magnitudes are comparable to those used for in vitro activation. Spatial distribution of MBV following intratumoural infusion was assessed by MRI and histology. **d**. Representative consecutive MRI cross sections overlaid with computed T_2_ relaxivity maps of the tumour. BV–*Pdl1* was used as the control. **e**. Representative histological sections of tumours treated with PBS or MBV–*Pdl1*. Tumour sections were stained for MNPs using Prussian blue and counterstained with nuclear red. The dashed red circle indicates the needle track. In vivo transduction was evaluated using MBV–*Pdl1* carrying an eGFP reporter. **f**. Flow cytometric quantification of eGFP^+^ cells in tumours treated with PBS or MBV–*Pdl1*. Tumours were divided into four fragments to illustrate intratumoural heterogeneity; statistical comparisons were performed using mouse-level means. **g**. Relative abundance of CD44□ tumour cells, ER-TR7□ fibroblasts, and CD45□ immune cells among eGFP□ cells for each treated mouse. PBS controls are not shown because eGFP□ events were at background levels. Assessment of off-target transduction and genome editing. **h**. Flow cytometric quantification of eGFP^+^ cells in major organs following PBS or MBV–*Pdl1* treatment. **i**. Next-generation sequencing (NGS) quantification of indel frequencies in tumours following PBS or MBV–*Pdl1* treatment. **j**. NGS quantification of indel frequencies in major organs following PBS or MBV–*Pdl1* treatment.

Magnetic nanoparticles enabled noninvasive tracking of MBV distribution by T□-weighted MRI in vivo (Supplementary Fig. S11). MRI and histological analyses revealed localized accumulation of MBV primarily surrounding the injection track (Fig. 4d, e), similar to other locally administered large nanoparticles ^31^. To quantify intratumoural transduction, tumours treated with MBV–*Pdl1*–eGFP were divided into four fragments and analyzed by flow cytometry. eGFP^+^ cells ranged from 0.6% to 2.7% across tumour fragments, reflecting heterogeneous intratumoural distribution typical of locally administered vectors within solid tumours (Fig. 4f). The majority of eGFP^+^ cells were CD44 tumour cells with lower levels detected in ER-TR7 fibroblasts, and CD45 immune cells (Fig. 4g). No eGFP expression was detected in major organs, and ex vivo fluorescence imaging further confirmed localization to treated tumours (Fig. 4h, Supplementary Fig. S12). Next-generation sequencing confirmed *Pdl1* indels in tumours but not in distal organs following MBV–*Pdl1* treatment (Fig. 4i, j). Importantly, the liver, which typically exhibits high off-target accumulation during treatment with conventional vectors, showed no detectable genome editing activity, despite the high transduction efficiency of BV in hepatocytes ^4,32^.

Consistent with functional confinement of genome editing, vector distribution was concentrated near the injection track, suggesting that editing occurred within spatially restricted regions of the tumour. Because tumours were dissociated without preserving spatial orientation relative to the injection track, both flow cytometric transduction estimates and bulk NGS measurements likely diluted locally edited regions with the substantially larger non-transduced tumour fraction. Despite this limited apparent coverage, MBV–*Pdl1* treatment produced robust therapeutic effects, indicating that localized checkpoint disruption within a restricted tumour niche can initiate tumour-wide immune remodelling.

Notably, tumour volumes diverged within three days of treatment (Fig. 5o), consistent with rapid immune-mediated cytotoxic effects that may reduce the abundance of initially transduced cells. In vitro, BV-mediated transgene expression in MC38 cells peaked at 24 h and declined over several days, becoming undetectable by day 7 (Fig. 2c), whereas prior in vivo studies demonstrated clearance of baculoviral transgene expression within 48 h in liver tissue ^4^. This contrast supports immune-mediated elimination of transduced cells in vivo, which would reduce detectable eGFP^+^ fractions in tumours.

**Fig. 5.**
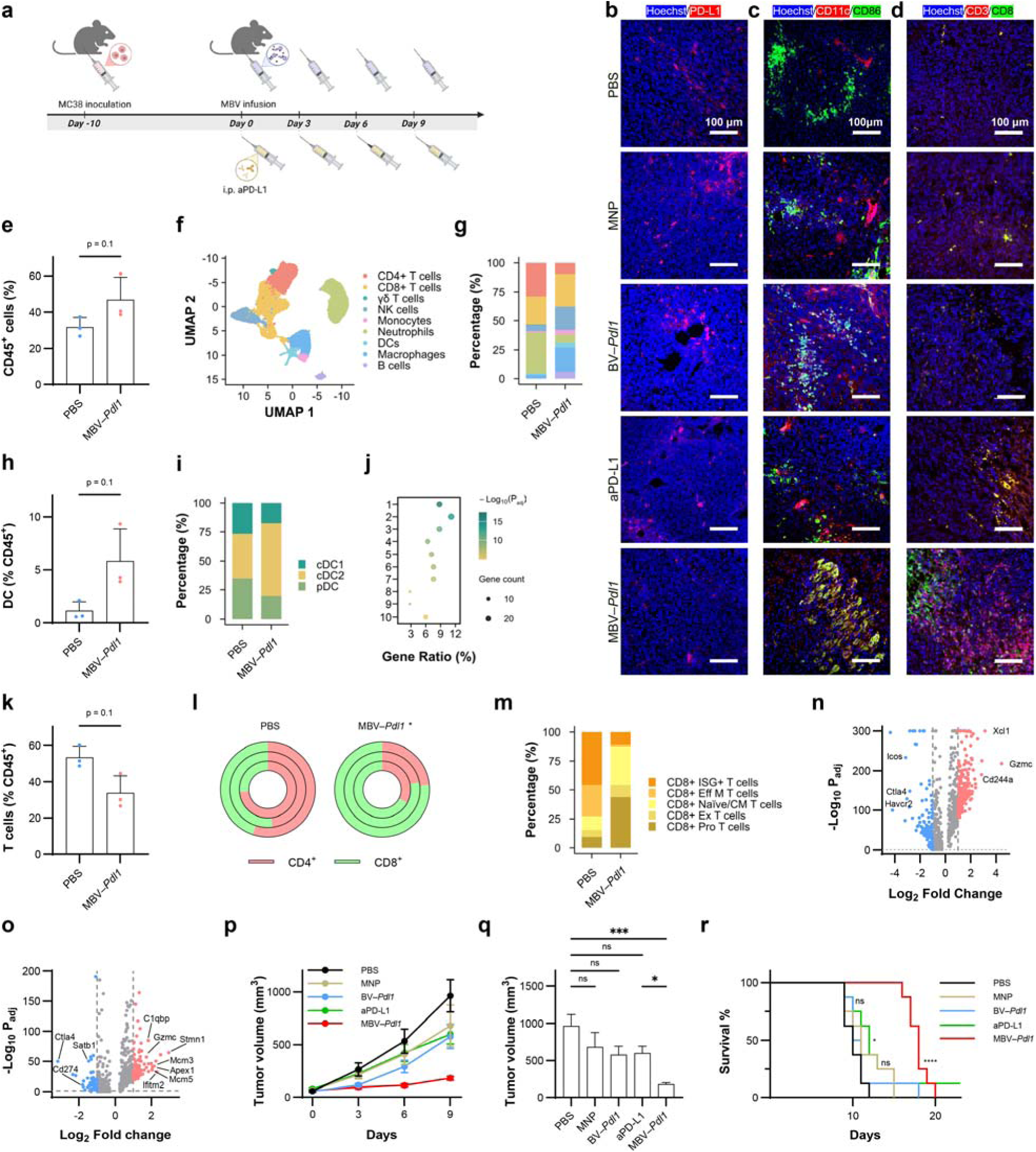
MBV–*Pdl1* modulates the tumour immune microenvironment and suppresses tumour growth. **a**. Schematic of the treatment timeline. **b-d**. Representative fluorescence microscopy images of tumours stained for PD-L1 (**b**), activated dendritic cells (CD11C^+^CD86^+^) (**c**), and CD8^+^ T cells (CD3^+^CD8^+^) (**d**) following treatment with PBS, MNP, BV–*Pdl1*, anti-PD-L1, or MBV–*Pdl1*. Nuclei were counterstained with Hoechst. **e**. Flow cytometry quantification of CD45^+^ cells in tumours treated with PBS or MBV–*Pdl1*. Cell population frequencies were compared at the mouse level using a two-sided Mann– Whitney U test (n = 3 mice per group). The effects of MBV–*Pdl1* treatment on the tumour immune microenvironment were further analyzed by single-cell RNA sequencing. **f**. UMAP embedding showing clustering of tumour-infiltrating immune cells. **g**. Proportions of major immune cell types in PBS- and MBV–*Pdl1*-treated tumours. **h**. Fraction of dendritic cells (DCs) among CD45^+^ cells in PBS or MBV–*Pdl1*-treated tumours. **i**. Proportions of DC subtypes in PBS- and MBV–*Pdl1*-treated tumours. **j**. Pathway enrichment analysis of DCs in MBV–*Pdl1*-treated tumour relative to PBS. Enriched pathways include: 1. Antigen processing and presentation; 2. Response to virus; 3. Activation of innate immune response; 4. Leukocyte mediated cytotoxicity; 5. Cell killing; 6. Positive regulation of cell-cell adhesion; 7. Regulation of T cell activation; 8. NADH metabolic process; 9. Glucose catabolic process; and 10. Lymphocyte proliferation. **k**. Fraction of T (γδ, CD4^+^, and CD8^+^) cells among CD45^+^ cells in PBS or MBV–*Pdl1*-treated tumours. **l**. Proportions of CD4^+^ and CD8^+^ T cells in PBS and MBV–*Pdl1*-treated tumours; each circle represents one mouse. **m**. Proportions of CD8^+^ T cell subtypes in PBS or MBV–*Pdl1*-treated tumours. **n**. Volcano plot of differentially expressed genes (|log_2_FC| > 1, *P*_adj_ < 0.05) in CD8^+^ T cells from MBV–*Pdl1*-tumours relative to PBS. **o**. Volcano plot of differentially expressed genes (|log_2_FC| > 1, *P*_adj_ < 0.05) in NK cells from MBV–*Pdl1*-tumours relative to PBS. Additional analyses are provided in Supplementary Fig. S15. Therapeutic efficacy of MBV–*Pdl1*. **p**. Tumour growth curves following the indicated treatment. Data are presented as mean ± s.e.m. **q**. Tumour volumes on day 9 after treatment initiation. Tumour volumes were compared using the Kruskal-Wallis test followed by Dunn’s multiple-comparisons test. Data are presented as mean ± s.e.m. ns, *, and *** denote no significance, *P* < 0.05, and *P* < 0.0001, respectively. **r**. Kaplan-Meier survival curves following the indicated treatment (n = 8 per group). ns, *, and **** denote no significance, *P* < 0.05, and *P* < 0.0001, respectively.

Although eGFP signal was detected in CD45^+^ immune cells in vivo, baculoviral transduction of hematopoietic-lineage cells is negligible under in vitro conditions ^33^. The presence of eGFP^+^ CD45^+^ cells is therefore most consistent with phagocytic uptake of transduced tumour cells or antigen transfer rather than productive viral transduction, consistent with enhanced recruitment and activation of antigen-presenting cells in treated tumours. Together, these observations indicate that the measured eGFP expression and indel frequencies underestimate peak vector engagement and editing activity within tumours.

Collectively, these results demonstrate that magnetic gating enables localized genome editing under complement-mediated constraint, resulting in functional confinement of genome editing activity to tumours while limiting systemic dissemination.

## MBV-mediated disruption of *Pdl1* prevents compensatory checkpoint signalling, enhances immune infiltration, and suppress tumour growth

Having established functional confinement of genome editing in vivo, we next examined how localized *Pdl1* disruption modulates the tumour immune microenvironment and therapeutic response. Therapeutic efficacy was evaluated by comparing PBS, MNP alone, BV–*Pdl1*, MBV–*Pdl1*, and systemic anti-PD-L1 antibody (aPD-L1) (Fig. 5a). Immunostaining confirmed reduced PD-L1 expression in MBV–*Pdl1*-treated tumours but not in BV–*Pdl1*-treated tumours (Fig. 5b). Although viral transduction was spatially heterogeneous, immunostaining revealed reduced PD-L1 expression across much of the tumour following MBV–*Pdl1* treatment. This observation is consistent with immune-mediated propagation of checkpoint modulation initiated by localized genome editing and antiviral sensing, which can extend beyond initially transduced cells and remodel the tumour microenvironment ^34,35,36^. Both BV–*Pdl1* and MBV–*Pdl1* treatments increased infiltration of CD11c^+^CD86^+^ dendritic cells (Fig. 5c), whereas MBV–*Pdl1* and aPD-L1 treatment increased CD3^+^CD8^+^ T cell infiltration (Fig. 5d).

To comprehensively characterize the impact of MBV–*Pdl1* on the tumour immune microenvironment, we analyzed CD45 immune cells isolated from tumours treated with PBS or MBV–*Pdl1*. Flow cytometry analysis revealed a near doubling of CD45 immune cells in MBV–*Pdl1*-treated tumours relative to PBS controls (Fig. 5e). Single-cell RNA sequencing of tumour-infiltrating immune cells demonstrated a marked shift in immune composition following MBV–*Pdl1* treatment (Fig. 5f, g). Tumours treated with PBS were dominated by three major populations – neutrophils, CD4^+^ T cells, and CD8^+^ T cells. MBV–*Pdl1* treatment increased immune diversity, with elevated proportions of macrophages, dendritic cells, B cells, natural killer (NK) cells, accompanied by reduced fractions of neutrophils, and CD4^+^ T cells.

Antigen-presenting cells, including macrophages, dendritic cells, and B cells, were substantially enriched following MBV–*Pdl1* treatment (Fig. 5g). Dendritic cells increased more than fourfold among CD45^+^ cells (Fig. 5h), with conventional type 2 dendritic cells (cDC2s) comprising the dominant subset (Fig. 5i, Supplementary Fig. S15a, b). Gene-expression analysis of tumour-infiltrating dendritic cells revealed enrichment of pathways associated with antiviral defense and antitumour immunity, including antigen processing and presentation, activation of innate immune responses, regulation of T cell activation, and type I interferon production (Fig. 5j).

Although the overall proportion of T cells decreased slightly, the CD8^+^/CD4^+^ T cell ratio increased significantly in MBV–*Pdl1*-treated tumours (Fig. 5k, l). Subclustering of CD8□ T cells further revealed a shift from effector and exhausted phenotypes toward naïve and cycling states (Fig. 5m, Supplementary Fig. S15c, d). PBS-treated tumours contained higher proportions of effector and exhausted CD8□ T cells, whereas MBV–*Pdl1*-treated tumours exhibited a marked increase in naïve and cycling CD8□ T cells. Differential expression analysis revealed upregulation of genes associated with proliferation and cytotoxic potential, including *Xcl1, Gzmc,* and *Cd244a*, alongside reduced expression of inhibitory receptors (*Ctla4*, *Icos*, and *Havcr2*) (Fig. 5n). CD4^+^ T cell analysis revealed that MBV–*Pdl1* treatment increased memory and regulatory T cell fractions (Supplementary Fig. S15e-g). NK cells were also enriched following MBV–*Pdl1* treatment (Fig. 5g). Analysis of exhaustion and inhibitory markers (*Ctla4*, *Cd274*, and *Satb1*) alongside proliferation-associated genes (*Stmn1*, *Mcm3*, *Mcm5*) and activation-associated genes (*Gzmc*, *Ifitm2*, *Apex1*, *C1qbp*) revealed that NK cells in MBV–*Pdl1*-treated tumours exhibited transcriptional signatures consistent with antiviral activation and cytotoxic effector programs (Fig. 5o).

MBV–*Pdl1* was administered every three days to account for heterogeneous intratumoural distribution and transient transgene expression, consistent with localized vector engagement (Fig. 5a). MBV–*Pdl1* produced the strongest tumour growth suppression among all groups, including aPD-L1, whereas MNP or BV–*Pdl1* alone induced only modest delay (Fig. 5p, q). MBV–*Pdl1* significantly prolonged survival, with a median survival of 18 days compared with 10 days for PBS and 12 days for aPD-L1 (Fig. 5r). No significant body weight loss was observed (Supplementary Fig. S16a).

Dose escalation of MBV–*Pdl1* did not further improve survival relative to standard dosing but remained superior to MBV–eGFP (Supplementary Fig. S16b, c), indicating that antiviral immune activation alone does not account for the therapeutic efficacy. Tumours derived from MC38 PD-L1 knockout cells exhibited only marginally reduced growth compared with wild-type tumours (Supplementary Fig. S16d-f), demonstrating that tumour-intrinsic PD-L1 loss alone is insufficient. Together, these findings indicate that therapeutic efficacy arises from the coordinated integration of antiviral immune activation and spatially confined *Pdl1* disruption, rather than either mechanism in isolation.

## Localized MBV–*Pdl1* activation synergizes with systemic CTLA-4 blockade

To assess compatibility with systemic immune checkpoint blockade, MBV–*Pdl1* was combined with anti-CTLA-4 antibody (aCTLA-4) in MC38 tumour-bearing mice (Fig. 6a). CTLA-4 blockade is a clinically relevant partner for PD-1/PD-L1-targeted therapies and complements localized checkpoint disruption by reducing regulatory T cell-mediated immunosuppression ^37^. The combination produced greater tumour suppression than either monotherapy or antibody combinations (Fig. 6b-f). Median survival increased to 38 days in the MBV–*Pdl1* + aCTLA-4 group compared with 10 days for PBS, 13.5 days for aCTLA-4 alone, and 14 days for aCTLA-4 + aPD-L1 (Fig. 6g). Two mice in the combination group achieved complete tumour regression. No significant body-weight loss was observed, indicating favorable tolerability (Supplementary Fig. S17a). In a pilot rechallenge experiment, the two long-term survivors from the MBV–*Pdl1* + anti-CTLA-4 group were re-implanted with MC38 cells and monitored for 20 days. Neither mouse developed detectable tumours (Supplementary Fig. S17b), consistent with durable antitumour immune protection.

**Fig. 6.**
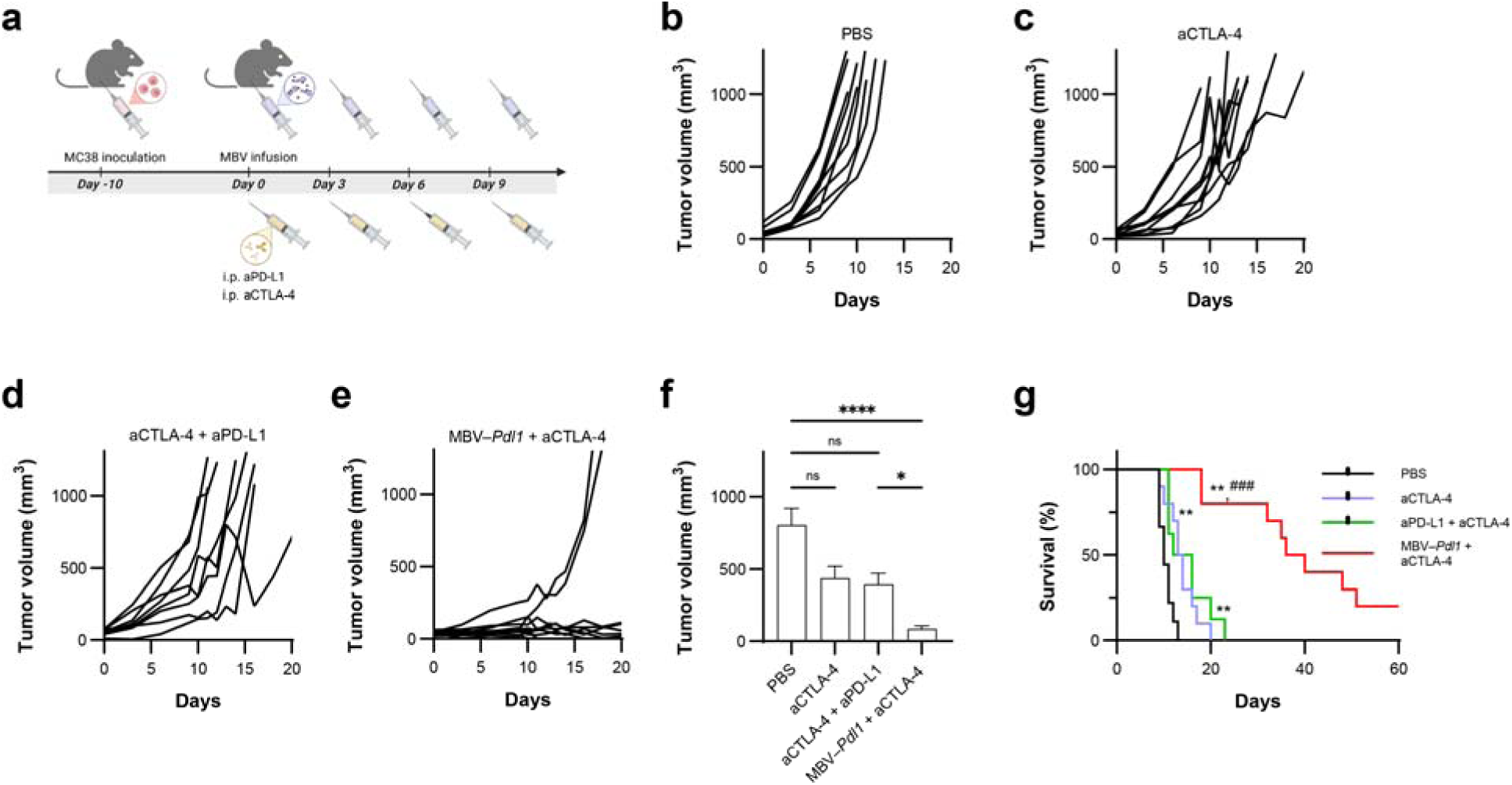
MBV–*Pdl1* and aCTLA-4 synergistically suppress tumour growth. **a**. Schematic of the treatment timeline. **b-e**. Tumour volumes measured from treatment initiation. Data are presented as mean ± s.e.m. **f**. Tumour volumes measured on day 9 after treatment initiation. Tumour volumes were compared across groups using the Kruskal-Wallis test followed by Dunn’s multiple-comparisons test. ns, *, and **** denote no significance, *P* < 0.05, and *P* < 0.0001, respectively. **g**. Corresponding Kaplan-Meier survival curves following the indicated treatment. ** denotes *P* < 0.01 versus PBS; ^###^ denotes *P* < 0.001 versus anti-PD-L1 + anti-CTLA-4.

## Discussion

In vivo genome editing provides a programmable approach to modulating disease-associated genes. In particular, CRISPR genome editing has become a fundamental tool for interrogating oncogenic drivers in cancer research. However, its clinical translation has been limited by the lack of strategies to precisely control genome editing activity within defined spatial contexts in vivo. Systemic dissemination of vectors risks unintended editing outside the target region. Previous studies have developed light- or chemical-inducible Cas9 nucleases ^38,39^; however, optical signals cannot penetrate deep tissue owing to scattering and absorption, and chemical activation remains constrained by the biodistribution of both Cas9 nucleases and activating agents. These approaches do not decouple genome editing activity from vector distribution and therefore cannot impose spatial control over editing activity.

Here, we integrate external magnetic regulation with intrinsic complement inactivation to achieve spatially gated control of genome editing activity in vivo. Local viral delivery has been demonstrated across diverse anatomical sites ^40^, and clinical studies of magnetically guided nanoparticle targeting have demonstrated localization of therapeutics in deep tissues, supporting the translational feasibility of magnetically gated activation. Although single-site intratumoural infusion produced localized vector deposition, broader coverage could be increased by multi-site delivery if needed. Such an approach is clinically plausible because spatially distributed placement is already used in procedures such as brachytherapy. Nonetheless, clinical deployment will require optimization of injection strategies and magnetic field geometries.

We developed a hybrid genome-editing platform that decouples genome editing activity from vector biodistribution by integrating complementary physical and biological constraints. Host complement imposes systemic restriction, whereas magnetic activation restores localized viral entry at the target site, establishing orthogonal biological and physical control of genome editing. Together, these constraints confine CRISPR activity to targeted tumour regions while limiting dissemination and unnecessary nuclease exposure. Although baculoviral transgene expression is transient, CRISPR-mediated editing irreversibly modifies the target locus, enabling sustained checkpoint disruption within the targeted tumour region without prolonged vector persistence.

Viral vectors, including oncolytic viruses, have long been leveraged in cancer immunotherapy owing to their capacity to stimulate innate and adaptive immunity. However, antiviral sensing can induce compensatory checkpoint upregulation, creating a regulatory feedback loop that restrains effector function and often necessitates systemic blockade. These observations highlight a key limitation of existing approaches, in which immune activation and regulatory feedback are not spatially coordinated. Our strategy addresses this limitation by embedding programmable checkpoint disruption within a spatially gated, non-replicative viral platform that restricts genome editing activity to defined tumour regions. Unlike replicating oncolytic viruses, MBV enables localized genome editing without sustained systemic vector persistence, thereby coupling immune activation and checkpoint disruption within the same tumour region while avoiding the safety and regulatory constraints associated with replication-competent genome-editing vectors.

Within tumours, MBV preserves innate immune activation while restricting genome editing activity to the activated field. Baculoviral transduction promotes antigen presentation and immune cell recruitment but also induces checkpoint upregulation. Local disruption of *Pdl1* resolves this inhibitory feedback while preserving the immunostimulatory context generated by viral sensing. The reduction of PD-L1 expression across much of the tumour despite heterogeneous viral distribution suggests that localized genome editing activity can initiate immune-mediated remodelling beyond the initially transduced cells. Consistent with this interpretation, the limited efficacy of tumour-intrinsic PD-L1 knockout and the modest response to systemic PD-L1 blockade indicate that therapeutic benefit arises from the coordinated integration of antiviral immune activation and spatially confined genome editing activity, rather than checkpoint disruption alone.

These findings suggest that uniform tumour-wide genome editing may not be required for therapeutic efficacy. Instead, checkpoint disruption within a restricted tumour region may be sufficient to initiate broader immune remodelling, consistent with a model in which localized antiviral activation and checkpoint relief generate a propagating antitumour response across non-edited tumour regions. This further supports a framework in which control of genome editing activity, rather than uniform delivery, governs therapeutic outcome.

At the tissue level, MBV–*Pdl1* increases immune cell density and diversification, particularly among antigen-presenting populations. Expansion of dendritic cells and enrichment of antigen-processing pathways are consistent with enhanced T-cell priming within tumours. Transcriptional profiling supports recruitment and activation of proliferative and cytotoxic effector populations rather than solely reinvigoration of pre-existing exhausted cells, suggesting durable immune reprogramming following localized genome editing. Spatial confinement focuses antiviral immune amplification within tumours while limiting detectable editing and vector activity in distal organs. Synergy with systemic CTLA-4 blockade further illustrates how localized checkpoint editing can integrate with systemic immunomodulation across spatial scales without overt toxicity. Although limited by sample size, rejection of MC38 rechallenge in two long-term survivors from the MBV–*Pdl1* plus anti-CTLA-4 group is consistent with the establishment of protective antitumour immunity and warrants further evaluation of durable immune memory.

Building on prior magnetic control of viral delivery, this study advances that framework by establishing functional control over therapeutic genome editing activity in vivo. Whereas our prior work established spatial confinement of editing, the present study demonstrates that physical regulation can be architecturally coupled to endogenous immune signalling to resolve tumour-intrinsic regulatory feedback. This approach further suggests that intrinsic immune clearance mechanisms, such as complement inactivation, can be harnessed as design features to actively constrain genome editing activity in vivo, providing a strategy for engineering next-generation vectors with built-in biological containment. The modular architecture of baculovirus permits incorporation of additional guide RNAs, enabling multiplexed and adaptable genome editing within spatially defined tissue regions.

Together, these findings establish MBV as a controllable genome-editing framework that integrates spatial regulation with endogenous immune activation to enable localized combination immunotherapy. By prioritizing control of genome editing activity over passive biodistribution, this strategy enables tumour-restricted immunomodulation while allowing localized genome editing events to initiate broader immune remodelling within the tumour microenvironment. More broadly, this study illustrates how external physical inputs can be integrated with intrinsic biological regulatory circuits to architect spatially controlled genome editing activity in vivo, establishing a generalizable framework for coupling physical and biological regulation in therapeutic systems.

## Online methods

### Materials

Iron(III) acetylacetonate (Fe(acac)3, 99.9%), oleic acid (90%), oleylamine (70%), 1,2-tetradecanediol (90%), benzyl ether (99%), chloroform (99%), toluene (99.9%), DMSO (99.9%), ferrozine (97%), were purchased from Sigma-Aldrich. 1,2-Distearoyl-sn-glycero-3-phosphoethanolamine-N-[methoxy (polyethylene glycol)-2000] (DSPE-mPEG_2000_) and 1,2-distearoyl-sn-glycero-3-phosphoethanolamine-N-[maleimide(polyethylene glycol)-2000] (DSPE-PEG_2000_-maleimide) were purchased from Avanti Polar Lipids. Cysteine-terminated TAT peptides (CGYGRKKRRQRRR) were synthesized by Genscript.

Bac-to-Bac baculovirus expression system and Cellfectin II was purchased from Thermo Fisher Scientific. BacPAK qPCR titration kit, NucleoMag® NGS Clean-up and Size Select kit, PrimeScript RT Master Kit, and TB Green Premix Ex Taq Master Mix were purchased from Takara Bio. Plasmids pX330 (catalog no. 42230), pCMV-GFP (catalog no. 11153), and pCMV-iRFP (catalog no. 45457) were obtained from Addgene. All gRNAs (Supplementary Table S1) and PCR primers (Supplementary Table S2, S3, and S4) were synthesized by Eurofins. xGen™ UDI Primers for NGS library preparation were purchased from Integrated DNA Technologies. Quick-DNA^TM^ MicroPrep kit and Quick-RNA Miniprep Kit were purchased from ZYMO Research. jetOPTIMUS® transfection reagent was purchased from Polyplus.

Enzymes, buffers, and kits used for molecular cloning, including Q5® Hot Start High-Fidelity 2× Master Mix, PCR & DNA Cleanup Kit, DNA Gel Extraction Kit, Quick CIP, T4 Polynucleotide Kinase, T4 DNA Ligase Reaction Buffer, BbsI-HF, T7 endonuclease I, and Quick Ligation Kit, were purchased from New England Biolabs. 10-beta competent E. coli cells were also obtained from New England Biolabs.

All antibodies are listed in Supplementary Table S5.

### Production of baculoviral vectors

Recombinant baculovirus vectors were generated using the Bac-to-Bac baculovirus expression system. Briefly, the expression cassettes of Cas9, gRNA, and reporter genes were inserted into the pFastBac vector and transformed into DH10Bac competent cells (Supplementary Fig. S2). The recombinant bacmids containing virus genome and the expression cassettes were extracted using the PureLink HiPure Plasmid Miniprep Kit and transfected into Sf9 insect cells using Cellfectin II according to the manufacturer’s protocol. The insect cell culture medium containing the budded viruses was centrifuged and filtered to remove cell debris using Bottle Top filters (0.45 μm; Thermo Fisher Scientific). The collected recombinant baculoviral vector was amplified in Sf9 cells for two more passages. Baculoviral stocks at passage 3 was used for all experiments. Viral particles were concentrated and purified by filtration through a 0.45 μm Bottle Top filter followed by centrifugation at 6,000 g for 3 h at 4 °C. The resulting viral pellets were resuspended in PBS and stored at 4 °C until use. Baculoviral titers were quantified using the BacPAK qPCR titration kit.

### Synthesis of magnetic nanoparticles

Magnetic nanoparticles (MNPs) were synthesized using a two-step method adapted from previously reported methods ^27^. Briefly, magnetite (Fe_3_O_4_) nanocrystals were prepared by thermal decomposition of iron(III) acetylacetonate (Fe(acac)_3_) in benzyl ether in the presence of oleic acid, oleylamine, and 1,2-hexadecanediol. To render the nanocrystals water-dispersible, they were coated with a mixture of DSPE-mPEG_2000_ and DSPE-PEG_2000_-maleimide at a molar ratio of 19:1 using a dual-solvent exchange method ^41^. For peptide conjugation, freshly coated MNPs were incubated with cysteine-terminated TAT peptides at a molar ratio of 1:200 in 0.25× PBS overnight at room temperature. Unconjugated peptides were removed by ultracentrifugation at 80,000 g for 35 min at 4 °C. The nanoparticles were washed, sterilized by filtration through a 0.22 μm syringe filter, and stored at 4 °C until use.

Nanocrystal size and size distribution of coated MNPs were characterized by transmission electron microscopy (TEM) and dynamic light scattering (DLS). Magnetization curve of nanocrystals was measured at 300K using a superconducting quantum interference device (Quantum Design MPMS). Particle concentrations were quantified using a Ferrozine assay. Successful surface conjugation of TAT peptides was confirmed by gel-shift assay (Supplementary Fig. S1).

### Cell culture

MC38 murine colon adenocarcinoma cells were obtained from Kerafast, and Sf9 insect cells were obtained from Thermo Fisher Scientific. All cell lines were cultured according to the manufacturers’ recommended protocols.

### In vitro BV transduction

To evaluate the efficiency and temporal dynamics of BV transduction, MC38 cells were seeded at a density of 4 x 10^4^ cells per well in 24-well plates and allowed to adhere overnight. Cells were then incubated with DMEM/F12 containing BV–eGFP at varying multiplicities of infection (MOIs) for 24 h, after which the medium was replaced with fresh DMEM/F12. eGFP expression in MC38 cells was quantified by flow cytometry at the indicated time points.

To assess magnetic activation and complement-mediated inhibition of BV transduction, MC38 cells seeded in 24-well plates were incubated with BV–eGFP in DMEM/F12, BV–eGFP in DMEM/F12 supplemented with 50% adult mouse serum (AMS), or MBV–eGFP in DMEM/F12 supplemented with 50% AMS. For cells treated with MBV–eGFP, culture plates were placed on a custom-designed magnetic plate for 30 min to enable magnetic activation (Supplementary Fig. S3). Cells were subsequently washed and maintained in DMEM/F12. eGFP expression in MC38 cells was assessed at 24 h by fluorescence microscopy and flow cytometry. To evaluate magnetic activation of *Pdl1* disruption under complement pressure, cells were treated with BV–*Pdl1* or MBV–*Pdl1* as described above. eGFP and PD-L1 expression were quantified by flow cytometry at 24 and 48 h post-treatment, respectively.

### Design of sgRNAs for *Pdl1* disruption

Six candidate single-guide RNAs (sgRNAs) targeting early exons or key structural regions of PD-L1 were designed using a bioinformatics pipeline incorporating the Vienna Bioactivity CRISPR score (Supplementary Table S1) ^28^. Potential on- and off-target sites were predicted using the CRISPRO web tool (http://crispor.gi.ucsc.edu/crispor.py) ^42^, and the top four predicted off-target loci for each sgRNA were selected for downstream next-generation sequencing (NGS) analysis.

MC38 cells were transfected with pX330-sgRNA plasmid using the jetOPTIMUS transfection reagent. Genomic DNA was harvested 48 h post-transfection using the Quick-DNA^TM^ MicroPrep kit. Genomic regions encompassing CRISPR-Cas9 cleavage sites were amplified by PCR using Q5 Hot Start High-Fidelity 2× Master Mix under the following conditions: initial denaturation at 98 °C for 30 s; 34 cycles of 98 °C for 10 s, 68 °C for 15 s, and 72 °C for 30 s, followed by a final extension at 72°C for 2 min. For T7 endonuclease I (T7E1) assays, mismatched heteroduplex DNA was generated by denaturing and reannealing 1 μg of purified PCR product, followed by digestion with T7 endonuclease I at 37 °C for 1 h. Digestion products were resolved on 2% agarose gels. Gene-modification efficiency was calculated using the formula: 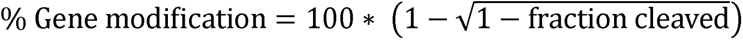.

For targeted NGS indel analysis, genomic regions spanning CRISPR-Cas9 cleavage sites were amplified by PCR. PCR amplicons were purified using the NucleoMag® NGS Clean-up and Size Select kit and subsequently indexed using xGen^TM^ UDI 10-nt Primer Plates 1-4 (Integrated DNA Technologies) through a second PCR step (98 °C for 30 s; 30 cycles of 98 °C for 10 s, 64 °C for 15 s, and 72 °C for 30 s; final extension at 72 °C for 2 min). Indexed libraries were purified, verified by 1% agarose gel electrophoresis, and quantified using a Qubit Fluorometer (Thermo Fisher Scientific). Sequencing was performed on an Illumina MiSeq platform. Indel frequencies and editing profiles were analyzed using CRISPResso2 (http://crispresso2.Pinellolab.org) ^43^.

### In vitro transcriptomic analysis of BV transduction

MC38 cells were transduced with BV–*Pdl1*, MBV–Cas9, or MBV–*Pdl1* for 48 h. Total RNA was isolated using the Quick-RNA Miniprep Kit. Gene-expression profiling was performed using the nCounter® Mouse Tumor Signaling 360™ Panel. Data normalization, differential gene-expression analysis, and pathway enrichment were conducted using the nSolver Analysis Software (v4.0). *P* values in differential gene-expression analysis were adjusted using the Benjamini and Yekutieli method for multiple testing correction.

### In vitro characterization of BV-induced immune responses

BV-induced immune responses were evaluated using coculture systems consisting of BV-transduced MC38 cells with either bone marrow-derived dendritic cells (BMDCs) or splenocytes.

To generate BMDCs, bone marrow cells were harvested from female C57BL/6 mice and cultured in RPMI-1640 medium supplemented with 10% fetal bovine serum (FBS), 100 U/mL penicillin, 100 μg/mL streptomycin, 1000 U/mL granulocyte-macrophage colony-stimulating factor (GM-CSF), and 800 U/mL interleukin-4 (IL-4) for 3 days. MC38 cells were transduced with BV–*Pdl1* or BV–Cas9 for 24 h, after which they were seeded at a density of 2000 cells per well in 96-well plates and allowed to adhere overnight. BMDCs were then added to MC38 cells at a ratio of 2:1 (BMDC:MC38). LPS-treated BMDCs (100 ng/mL) were used as a positive control. After an additional 24 h of coculture, total RNA was extracted using the Quick-RNA Miniprep Kit. RNA was reverse transcribed into cDNA using the PrimeScript RT Master Kit. Gene expression was quantified by real-time quantitative PCR (RT-qPCR) using TB Green Premix Ex Taq Master Mix in a Bio-Rad CFX96 real-time PCR system. Relative gene expression was calculated using the 2^-ΔΔCt^ method.

In parallel experiments, splenocytes were isolated from the spleens of female C57BL/6 mice and cultured in RPMI-1640 medium supplemented with 10% FBS, 100 U/mL penicillin, and 100 μg/mL streptomycin. MC38 cells were transduced with BV for 24 h and subsequently seeded at 2000 cells per well in 96-well plates overnight. Splenocytes were then added at a ratio of 5:1 (splenocytes:MC38). LPS-treated splenocytes were used as a positive control. After 24 h of coculture, cytotoxicity was assessed using a Cell Counting Kit-8 (CCK-8) assay. Gene expression was quantified by RT-qPCR as described above.

### Design of the magnetic field for intratumoural MBV activation

The magnetic apparatus used for intratumoural MBV activation was designed based on magnet availability, tumour dimensions, and simulated magnetic force distribution. The magnetic force exerted on MNPs was calculated using the following equation ^44^:

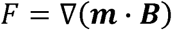

where *µ* denotes the magnetic permeability of the surrounding tissue, ***m*** is the magnetic moment of the MNPs, and ***B*** represents the magnetic flux density of the applied magnetic field.

Magnetic field distributions were computed by defining the magnet geometry and computing the spatial magnetic flux density generated by the defined magnet geometry, followed by calculation of magnetic force based on field gradients and a Langevin model of nanoparticle magnetization. Specifically, the computational workflow involved: (i) defining the geometry and spatial arrangement of permanent magnets, (ii) calculating the magnetic field distribution over a discretized spatial domain, (iii) estimating nanoparticle magnetization under the applied field using a Langevin function, and (iv) computing the resulting magnetic force from the field gradients. Spatial maps were generated over a discretized domain representing the tumour and surrounding tissues.

Based on the numerical simulation, four cylindrical NdFeB magnets (N52 grade; K&J Magnetics) were arranged in an alternating polarity configuration to maximize the magnetic field gradient (Supplementary Fig. S9). A custom 3D-printed animal bed was used to reproducibly position tumours at the center of the magnet array. The resulting magnetic field and force distributions were simulated using a MATLAB program developed in-house.

### Magnetically activated *Pdl1* disruption in vivo

All animal studies were approved by the Institutional Animal Care and Use Committee at the University of Kentucky. Female C57BL/6 mice (4-5 weeks old) were purchased from Charles River Laboratories. To establish the syngeneic mouse tumour model, mice received a subcutaneous injection of 5×10^5^ MC38 cells suspended in 100 μL of PBS into the right flank. Tumour volume and body weight were monitored every 3 days. Tumour volume was calculated using the formula V = (L × W^2^) / 2, where L and W denote tumour length and width, respectively. Treatments were initiated when tumours reached approximately 50 mm^3^. Mice were euthanized when tumour volume exceeded 1000 mm^3^ in accordance with human endpoint guidelines.

In monotherapy studies, mice were randomized into five groups receiving PBS, MNP alone, BV–*Pdl1* alone, MBV–*Pdl1*, or systemic anti-PD-L1 antibody (aPD-L1). MBV–*Pdl1* was prepared by mixing 1.5×10^7^ pfu of BV–*Pdl1* with 5 µg of MNP-TAT in 20 μL of PBS. Mice received intratumoural infusion of MBV–*Pdl1* every 3 days for a total of four doses. The injectate was delivered using a fanning technique, in which the needle was partially withdrawn and redirected to facilitate intratumoural distribution. The injection volume was fixed at 0.3 × the tumour volume while maintaining constant vector concentration, and the solutions were infused over approximately 5 min using a syringe pump. For mice treated with MNP alone or BV–*Pdl1* alone, injected doses were matched to the corresponding components in the MBV–*Pdl1* formulation. In the MNP alone or MBV–*Pdl1* groups, mice were positioned laterally on a 3D-printed animal bed immediately following infusion, with tumours aligned within the magnetic field and maintained under anesthesia for 1 h (Supplementary Fig. S9). Systemic aPD-L1 was administered intraperitoneally at a dose of 200 μg in 100 μL PBS on the same dosing schedule.

For combination therapy studies, mice were randomized into four groups receiving PBS, systemic aCTLA-4, systemic aCTLA-4 combined with aPD-L1, or MBV–*Pdl1* combined with systemic aCTLA-4, administered every 3 days for a total of four doses. In the aCTLA-4 group, mice received intraperitoneal injection of 200 μg (day 0) or 100 μg (day 3, 6 and 9) of aCTLA-4 in 100 μL PBS ^45^. In the aCTLA-4 plus aPD-L1 group, mice received intraperitoneal injection of aCTLA-4 as above and 100 μg aPD-L1 in 100 μL PBS per treatment. In the MBV–*Pdl1* plus aCTLA-4 group, MBV–*Pdl1* was administered intratumourally as described above, followed by intraperitoneal injection of aCTLA-4 as described above.

### Evaluation of in vivo distribution of MBV

To assess the intratumoural distribution of MBV following infusion, tumour-bearing mice received a single intratumoural injection of BV or MBV as described above. In vivo magnetic resonance imaging (MRI) was performed using a 7 T small animal MRI system (PharmaScan, Bruker) equipped with a 38-mm surface coil. Tumours were imaged using a spin-echo sequence with the following parameters: repetition time (TR) = 2,000 ms; echo time (TE) incremented from 12 ms to 144 ms in 12 ms steps; matrix size = 256 × 256; and field of view (FOV) = 30 mm. T_2_ relaxation maps were generated using a custom MATLAB script.

Following MRI, mice were euthanized, and tumours were harvested, cryosectioned, and stained with Prussian blue to visualize intratumoural MNP distribution.

To assess off-target activities of MBV–*Pdl1* in normal tissues, tumour-bearing mice were treated with PBS or MBV–*Pdl1*–eGFP. To minimize underestimation of off-target events resulting from immune clearance of edited cells, MBV–*Pdl1*–eGFP was administered once at a fourfold dose. 24 h after treatment, mice were euthanized and tumours as well as major organs (heart, kidney, lung, spleen, and liver) were harvested. Each tumour was cut into 4 fragments. Tumour fragments and organs were enzymatically dissociated using an enzyme cocktail (20 µg/mL DNase I, 2 mg/mL Collagenase Type IV, and 0.5 mg/mL Hyaluronidase Type 1-S). Cells were stained with APC-conjugated rat anti-mouse CD45 antibody, PE/Cyanine-conjugated rat anti-mouse/human CD44 antibody, Alexa Fluor 647-conjugated rat anti-mouse/rat/human ER-TR7 antibody and analyzed by flow cytometry.

To assess the *Pdl1* disruption efficiency in the whole tumour, 4 more tumours were harvested 24 h after a single injection of MBV–*Pdl1* and minced into 10-12 pieces. Genomic DNA was extracted from mechanically homogenized tissue with a Quick-DNA MicroPrep kit and subjected to targeted NGS amplicon sequencing.

### Immunofluorescence staining

To assess PD-L1 expression and dendritic cell and T-cell infiltration, mice were euthanized 3 days after completion of four treatment cycles, and tumours were harvested, fixed in 4% paraformaldehyde on ice for 2 h, and cryosectioned. Tissue sections were treated with 0.3% H_2_O_2_ for 10 min at room temperature before staining. For PD-L1 detection, sections were incubated with PE-conjugated rat anti-mouse PD-L1 antibody. For visualization of CD3^+^CD8^+^ T cells, sections were blocked with 5% goat serum for 1 h, and incubated with rabbit anti-mouse CD8α antibody for 1 h at room temperature, followed by incubation with Alexa Fluor 594-conjugated rat anti-mouse CD3 antibody and Alexa Fluor 488-conjugated goat anti-rabbit IgG antibody for an additional 1 h. Sections were counterstained with Hoechst 33342 for 5 min at room temperature and mounted using VectaShield Antifade Mounting Media (Vector Laboratories). Similarly, sections were stained for dendritic cells (CD11c^+^CD86^+^). Fluorescence images were acquired using a Nikon Eclipse Ti2 Inverted fluorescence microscope.

### Single-cell RNA sequencing

Immunological changes induced by MBV–*Pdl1* were characterized by single-cell RNA sequencing (scRNA-seq) of tumour-infiltrating immune cells isolated from 3 PBS-treated and 3 MBV–*Pdl1*-treated tumours. Two days after the fourth treatment, mice were euthanized and tumours were harvested for isolation of CD45^+^ cells following the 10x Genomics Cell Preparation protocol. Tumours were minced into 2-4 mm^3^ fragments and enzymatically dissociated using a tumour dissociation kit (Miltenyi Biotec) for 20 min at 37 °C. Cell suspensions were filtered through a 40-μm cell strainer and centrifuged at 500 g for 7 min at 4 °C and subjected to blood cell lysis. After washing, cells were labeled with APC-conjugated anti-mouse CD45 antibody and sorted using a BD FACSymphony^TM^ S6 Cell Sorter (Supplementary Fig. S13). For each tumour, up to 10,000 CD45^+^ cells were loaded onto a Chromium Controller (10x Genomics) for single-cell encapsulation. Libraries were prepared according to the manufacturer’s instructions and sequenced on an Illumina HiSeq 2500 platform.

Raw sequencing data were processed using an R-based pipeline. Gene expression libraries were aligned to the mouse reference genome GRCm39 (10x Genomics pre-built reference, GRCm39-2024-A) using Cell Ranger (v8.0.1). Downstream analyses were performed with Seurat (v5.3.0) ^46^.

The computational workflow consisted of: (i) quality control filtering to remove low-quality cells and genes, (ii) normalization and scaling of gene expression data, (iii) dimensionality reduction using principal component analysis (PCA), (iv) batch correction and integration across samples using Harmony (v1.1.0) ^47^, (v) construction of a shared nearest neighbor (SNN) graph and unsupervised clustering using the Louvain algorithm, and (vi) visualization using Uniform Manifold Approximation and Projection (UMAP). Cell clusters were annotated based on canonical marker gene expression (Supplementary Fig. S14).

Quality control filtering was implemented as follows: cells with ≤200 detected genes and genes expressed in fewer than three cells were excluded; cells with total unique molecular identifier (UMI) counts or detected gene numbers below the 1st percentile or above the 99th percentile were removed; additional filtering excluded cells with ≤1,000 UMIs, ≤500 detected genes, or ≥5% mitochondrial transcript content. Putative doublets were identified and removed using scDblFinder (v1.17.0) ^48^. After filtering, a total of 30,006 cells (6,405 (mean) for PBS and 3,597 (mean) for MBV–*Pdl1*) were retained for analysis.

Major immune lineages, including T cells, macrophages, and dendritic cells, were subsetted for independent subclustering analysis. Within each subset, data were renormalized and re-clustered using the same Harmony-based workflow, and subclusters were annotated based on established lineage- and state-specific markers.

Differential gene expression analysis between clusters or experimental groups was performed using the Wilcoxon rank-sum test. Gene set enrichment analysis (GSEA) was conducted using the clusterProfiler package (v4.16.0) ^49^. Differences in cell-type proportions between experimental groups were assessed using a hypergeometric test. Comparisons of cell-state abundances across mice were considered exploratory due to limited biological replicates and were supported by orthogonal validation assays.

### Statistical analysis

All statistical analyses were performed using GraphPad Prism. Data were analyzed using two-sided unpaired t-tests (with Welch’s correction when variances were unequal), one-way analysis of variance (ANOVA) with post-hoc Dunnett’s test, or non-parametric tests (Mann–Whitney U test) as appropriate. Tumour volumes at day 9 post-treatment – the last time point at which all mice remained alive prior to control animals reaching humane endpoint criteria – were compared using the Kruskal-Wallis test followed by Dunn’s multiple-comparisons test. Survival data were analyzed using the Log-rank (Mantel-Cox) test. Exact p-values or significance thresholds and sample sizes are reported in the figure legends. Differential expression thresholds were defined as log fold change (|log FC|) > 0.58 for bulk RNA sequencing and |log FC| > 1 for single-cell RNA sequencing to account for differences in signal variability and noise characteristics between datasets.

## Schematic illustrations

Schematic illustrations were created with Biorender.com.

## Online content

Any methods, additional references, source data, supplementary information, acknowledgements, details of author contributions and competing interests; and statements of data and code availability are available online.

## Code Availability

Custom code used in this study is available from the corresponding author upon reasonable request. The codebase includes:

- MATLAB scripts for simulation of magnetic flux density and magnetic force distributions, including (i) spatial mapping at the tumor surface and (ii) whole-body magnetic force distribution in a mouse model.
- R scripts for processing, analysis, and visualization of single-cell RNA sequencing (scRNA-seq) data.

All MATLAB scripts were developed and tested using MATLAB R2024b. All R scripts were developed and tested using R version 4.4.2 within RStudio 2025.09.2 (Build 418).

Details of simulation parameters and analysis workflows are provided in the Methods section. Standard packages for scRNA-seq analysis (including Seurat and related visualization tools) were used as described.

## Data availability

The datasets generated and/or analyzed during this study are available from the corresponding author upon reasonable request. Sequencing data will be deposited in a public repository (e.g., GEO) upon publication.

## Acknowledgements

This work was supported by the NIH/NIBIB funding (R01EB026893 to S.T.). The authors would like to thank the U.K. Electron Microscopy Center supported by the National Science Foundation (NNCI-2025075), the U.K. Flow Cytometry & Immune Monitoring core facility supported by an NCI Center Core Support Grant (P30 CA177558), the OncoGenomics Shared Resource Facility of the University of Kentucky Markey Cancer Center (P30CA177558), and the U.K. Bioelectronics and Nanomedicine Research Center Core facility access award (to X.Y.).

## Author contributions

X.Y. and S.T. designed the project. X.Y., Z.Y., and S.T. performed the experiments and collected the data. X.Y., L.T., Y.P., J.H., D.H., J.L., C.W., and S.T. performed data analysis. X.Y. and S.T. wrote the original manuscript. X.Y., Y.P., Y.L., and S.T. revised the manuscript. Y.L. and S.T. supervised the project. All authors have given approval to the final version of the manuscript.

## Competing interests

X.Y. and S.T. are inventors on a pending, unlicensed patent application assigned to the University of Kentucky and related to the technologies described in this work.

**Correspondence and requests for materials** should be addressed to Sheng Tong.

