## Supplementary figures and tables for "Spatial control of genome editing activity enables localized immunotherapy"

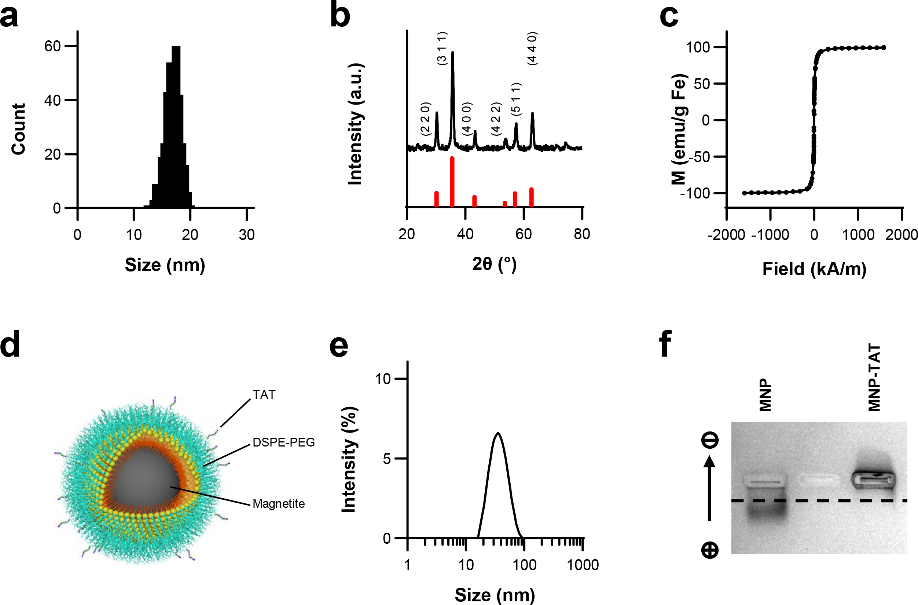

**Fig. S1 | Characterization of magnetic nanoparticle**.

**a**. Size distribution of iron oxide nanocrystals determined from transmission electron microscopy images (ImageJ analysis). The mean core diameter was 16.9 ± 1.3 nm. **b**. X-ray diffraction (XRD) pattern of iron oxide nanocrystals measured using an X-ray diffractometer (Bruker D8 ADVANCE). Red lines indicate the standard diffraction peaks of magnetite. The measured pattern is consistent with the characteristic cubic spinel structure of magnetite. **c**. Magnetization curves of magnetite nanocrystals measured at 300K using a superconducting quantum interference device (Quantum Design MPMS). **d**. Schematic of a magnetic nanoparticle (MNP) conjugated with TAT peptides. **e**. Hydrodynamic size distribution of magnetic nanoparticles measured by dynamic light scattering (DynaPro Nanostar, Wyatt Technology). The mean hydrodynamic diameter was 38.2 ± 0.2 nm with a polydispersity index of 0.231 ± 0.003. **f**. Agarose gel electrophoresis of MNPs with and without TAT peptide conjugation in Tris-acetate EDTA (TAE) buffer (pH 8.0). The dashed line indicates the well position. Maleimide-functionalized MNPs migrated toward the anode due to the negative charge of hydrolyzed maleimide (maleic acid), whereas MNP-TAT exhibited slight migration toward the cathode owing to the net positive charge of the conjugated peptides.

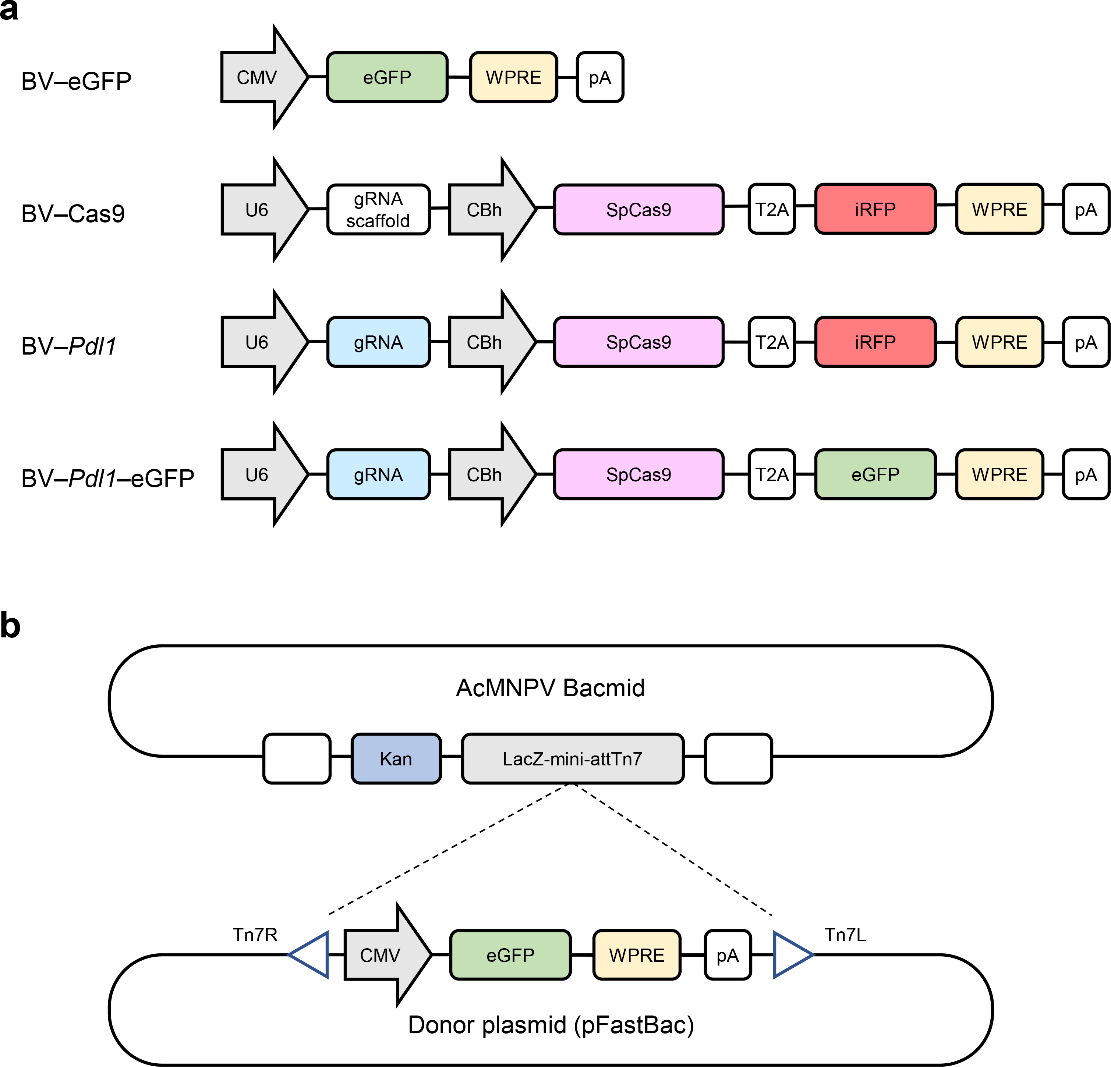

**Fig. S2 | BV plasmid design**.

Baculoviral vectors were generated through three sequential steps. First, expression cassettes encoding Cas9, guide RNAs, and reporter genes (**a**) were assembled using plasmids purchased from Addgene. Second, the expression cassettes were cloned into a donor plasmid (pFastBac), which was subsequently integrated into a bacmid by transformation into DH10Bac competent cells (**b**).

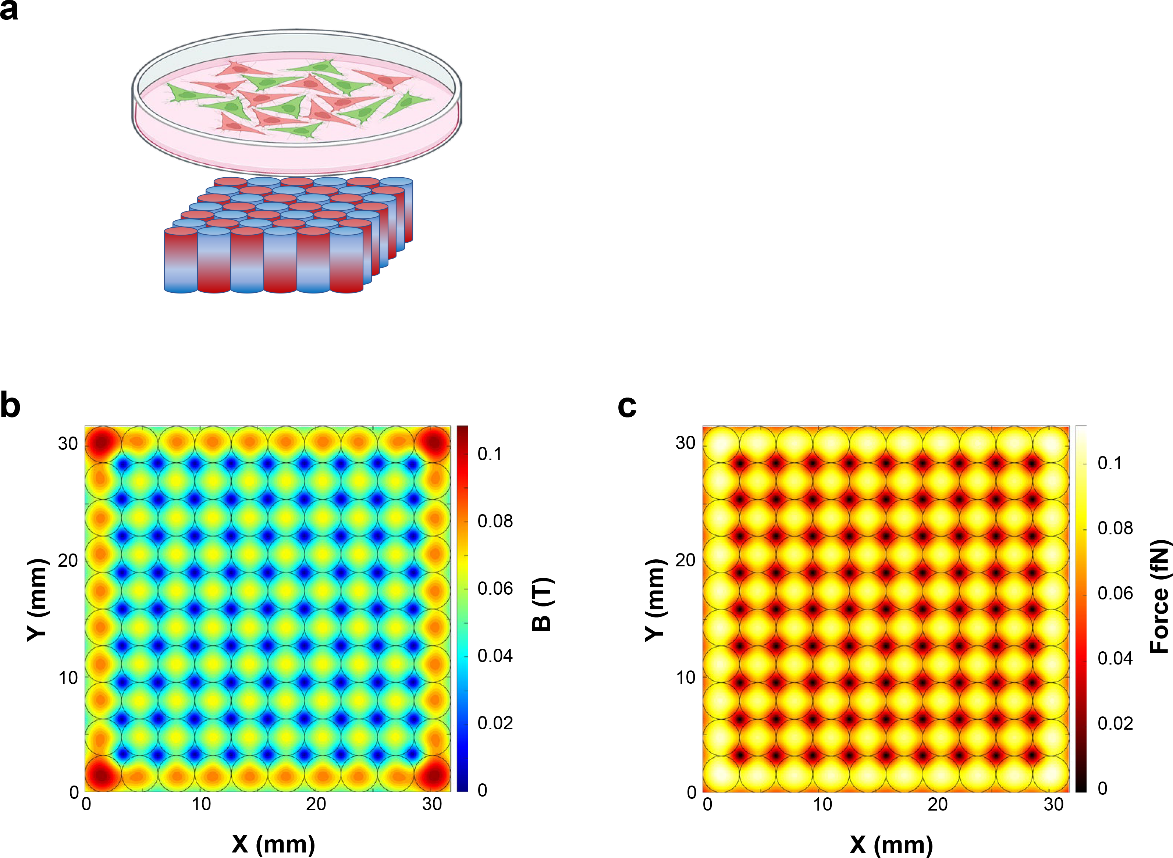

**Fig. S3 | Magnetic activation in vitro**.

**a**. Schematic of the in vitro magnetic activation setup. A matrix of magnets was placed beneath the cell culture Petri dish. The magnets were N52-grade NdFeB cylindrical magnets with dimensions of 1/8” × 1/4” (diameter × height). Red and blue colors denote the north and south magnetic poles, respectively. The distance between the top surface of the magnets and the cells in the Petri dish was approximately 2 mm. **b**, **c**. Contour plots of the magnetic flux density (**b**) and magnetic force (**c**) at the surface of the Petri dish, calculated for a 10 × 10 magnet matrix. A matrix larger than the culture surface was used in all experiments to avoid edge accumulation of MBV. Black circles denote magnet positions. The magnetic force was calculated for magnetite nanoparticles with a core diameter of 16.9 nm.

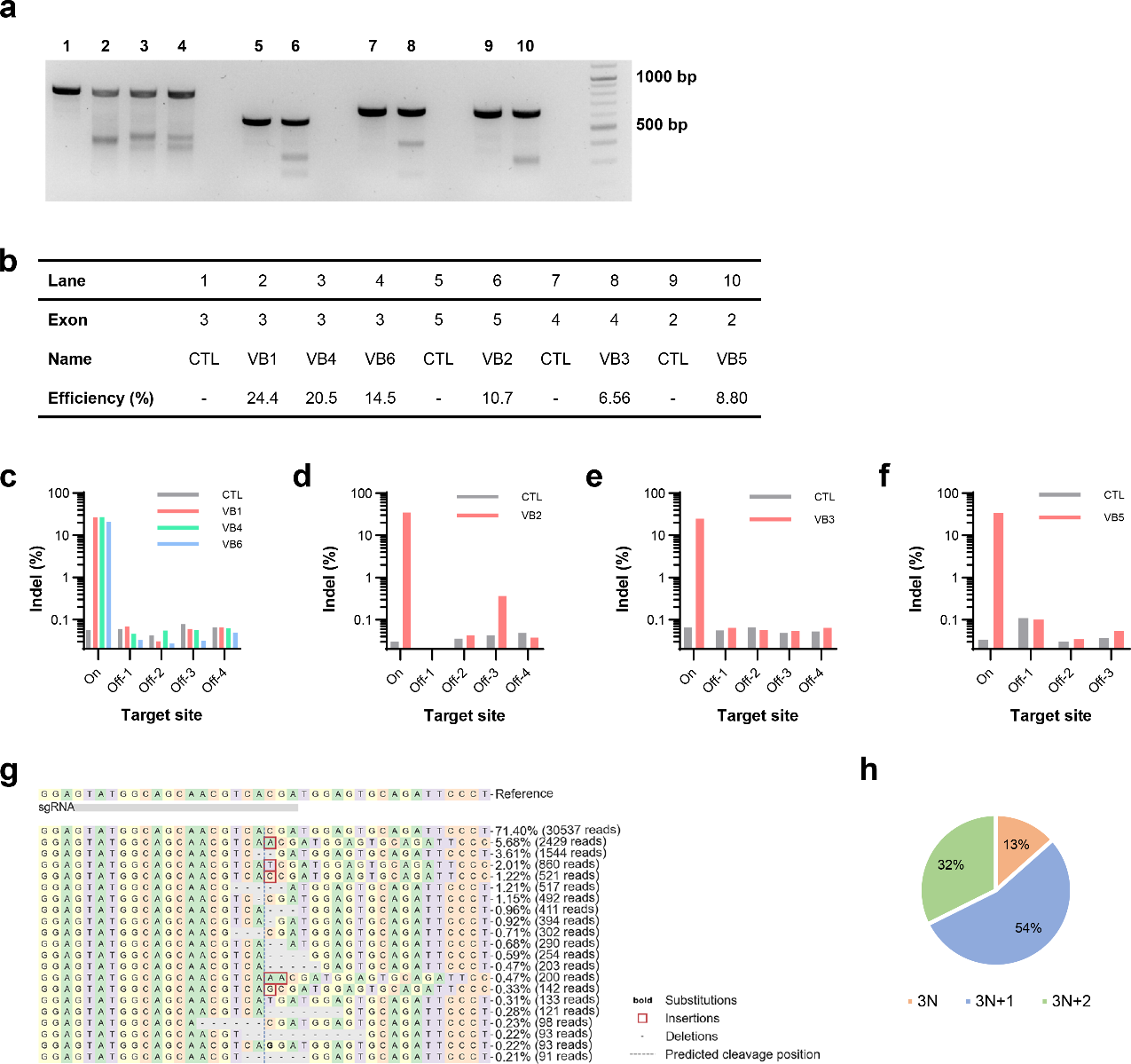

**Fig. S4 | gRNA screening**.

MC38 cells were transfected with pX330 plasmids encoding Cas9 and indicated guide RNAs. gRNA performance was evaluated by assessing on- and off-target efficiencies using the T7 endonuclease I (T7E1) assay and next generation sequencing (NGS), as described in the Methods. **a**. T7E1 analysis of all six candidate gRNAs. **b.** Indel efficiency quantified from the gel shown in **a**. **c**-**f**. Evaluation of on- and off-target activities of VB1, VB4, and VB6 (**c**), VB2 (**d**), VB3 (**e**), and VB5 (**f**), respectively, by NGS analysis. VB1 was selected based on its high on-target activity and minimal off-target effects. **g**. Representative VB1-induced mutation patterns quantified by NGS. **h**. Distribution of VB1-induced insertion and deletion (Indel) events categorized by resulting reading-frame shifts, quantified by NGS.

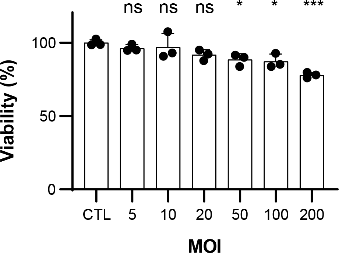

**Fig. S5 | Cytotoxicity of BV**.

To evaluate BV cytotoxicity, MC38 cells were seeded at a density of 2000 cells per well in 96 well plates and allowed to attach overnight. Cells were incubated with BV–*Pdl1* at the indicated multiplicities of infection (MOIs) for 24 hours. Cells were then cultured in fresh medium containing Cell Counting Kit-8 (CCK-8) for 3 h, and absorbance was measured at 450 nm using a microplate reader. Cytotoxicity was calculated as the ratio of absorbance relative to untreated control cells. Data are presented as mean ± s.d. ns, *, and *** denote no significance, *P* < 0.05, and *P* < 0.001 versus control, respectively.

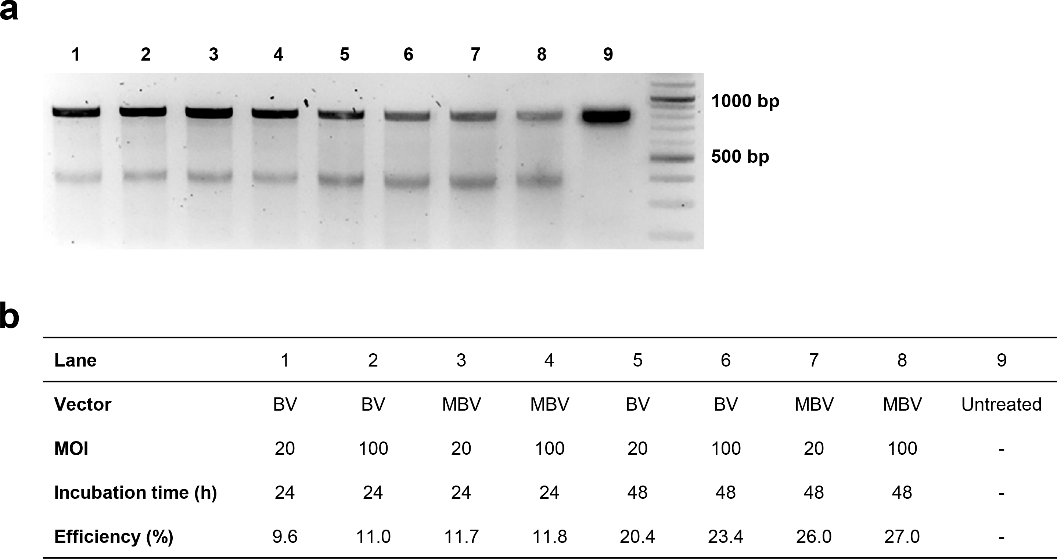

**Fig. S6 | Efficiency of BV- and MBV-mediated *Pdl1* disruption.**

MC38 cells were incubated with BV–*Pdl1* or MBV–*Pdl1* under the indicated conditions. Genomic DNA was harvested using the Quick-DNA^TM^ MicroPrep kit. Genomic regions spanning the CRISPR-Cas9 cleavage sites were amplified by PCR, and gene disruption efficiency was evaluated using the T7 endonuclease (T7E1) assay as described in the Methods. **a**. Gel electrophoresis of T7E1-digested PCR products. **b**. Incubation conditions and indel efficiencies quantified from the gel analysis. MBV showed higher *Pdl1* disruption efficiency relative to BV.

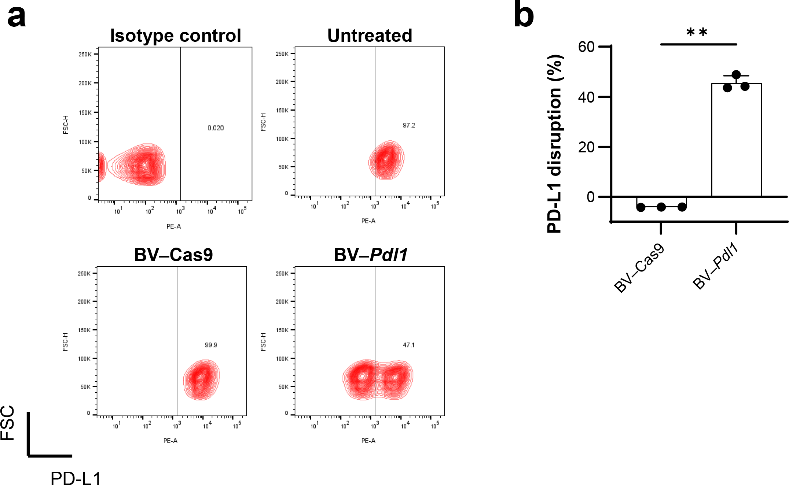

**Fig. S7 | Effect of BV–Cas9 and BV–*Pdl1* on PD-L1 expression**.

To assess the effect of BV–Cas9 and BV–*Pdl1* on PD-L1 expression, MC38 cells were incubated with BV–Cas9 or BV–*Pdl1* at a multiplicity of infection (MOI) of 20 for 24 h. Cells were then cultured in fresh medium for an additional 24 h, after which PD-L1 expression was quantified by flow cytometry. **a**. Representative flow cytometry contour plots. **b**. PD-L1 disruption efficiency calculated as the percentage of PD-L1-negative cells relative to untreated control cells. BV–Cas9 resulted in increased PD-L1 signal, whereas BV–*Pdl1* treatment produced a bimodal PD-L1 distribution, with distinct PD-L1-positive and -negative cell populations. Data are presented as mean ± s.d. ** denotes *P* < 0.01.

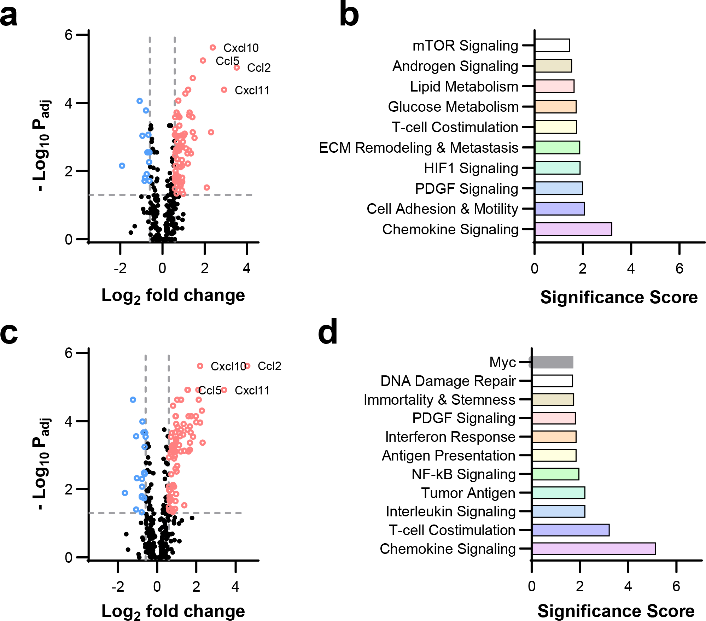

**Fig. S8 | Gene expression in MC38 cells treated with BV–*Pdl1.***

Transcriptomic responses of MC38 cells treated with BV–*Pdl1* and MBV–Cas9 were analyzed using the nCounter® Tumor Signaling 360 Panel as described in the Methods. All analyses were performed using nSolver Analysis Software (v4.0). **a**. Differential gene expression (|log_2_FC| > 0.58, *P*_adj_ < 0.05) in MC38 cells treated with BV–*Pdl1* relative to PBS control (n = 3 per group). *P* values were adjusted using the Benjamini and Yekutieli method for multiple testing correction. **b**. Pathway enrichment analysis of BV–*Pdl1*-treated MC38 cells relative to PBS control. **c**. Differential gene expression (|log_2_FC| > 0.58, *P*_adj_ < 0.05) in MC38 cells treated with MBV–Cas9 relative to PBS control (n = 3 per group). *P* values were adjusted using the Benjamini and Yekutieli method for multiple testing correction. **d**. Pathway enrichment analysis of MBV–Cas9-treated MC38 cells relative to PBS control.

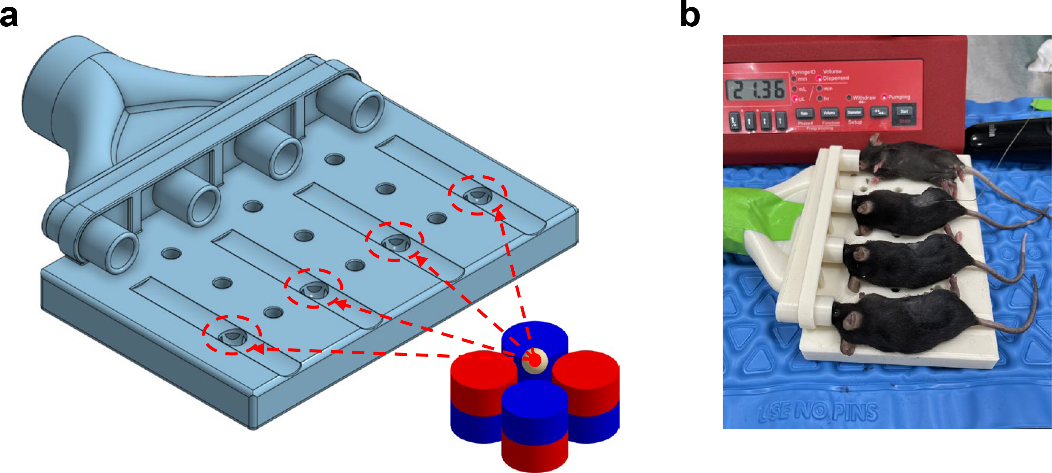

**Fig. S9 | Schematic of in vivo magnetic activation of MBV**.

**a**. Schematic of the custom-designed animal bed. Red dashed circles indicate flank tumor slots. An array of four magnets was installed beneath each tumor slot from the underside of the bed. The magnets were N52-grade NdFeB cylindrical magnets with dimensions of 1/2” × 1/2” (diameter × height). Red and blue colors denote the north and south magnetic poles. The dimensions and relative positions of the magnets and tumor are drawn to scale. **b**. Representative photograph of in vivo magnetic activation. From top to bottom, mouse 1 was undergoing magnetic activation, mouse 2 was receiving MBV infusion controlled by a syringe pump, and mice 3 and 4 were awaiting treatment.

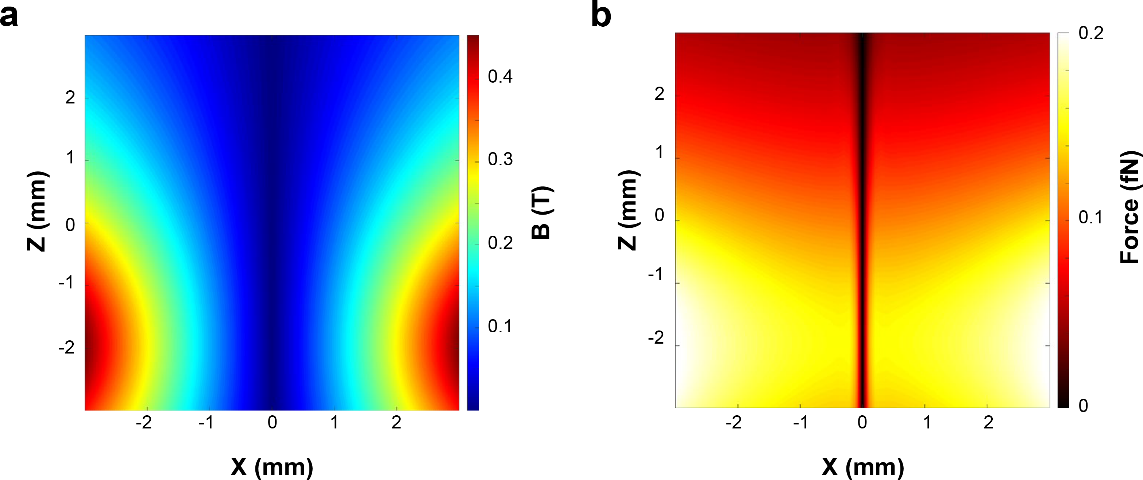

**Fig. S10 | Simulated magnetic fields on the tumor midplane**.

**a**, **b**. Contour plots of the magnetic flux density (**a**) and magnetic force (**b**) on the central plane of tumors. The geometry of the magnets and tumor is shown in Fig. 4**b**. The magnetic force was calculated for magnetic nanoparticles with a core diameter of 16.9 nm.

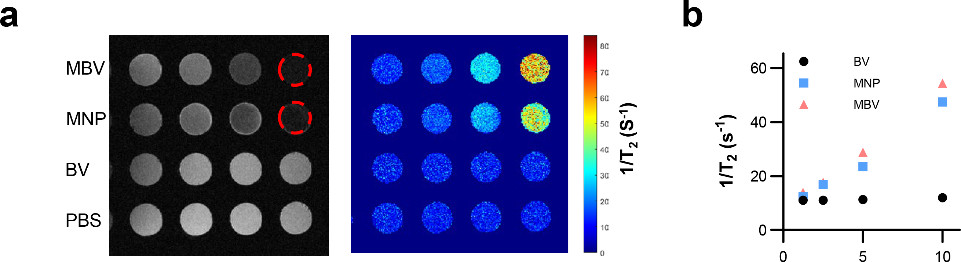

**Fig. S11 | MRI T_2_ relaxivity of MBV**.

To evaluate the T_2_ contrast properties of MBV, MBV was prepared as described in the Methods. Solutions of MNP, BV, and MBV was loaded into a 3D-printed mold and imaged using a 7 T small animal MRI system (PharmaScan, Bruker) equipped with a 38-mm surface coil. MNP and MBV solutions were matched for iron concentration, and BV and MBV solutions were matched for BV concentration. **a**. T_2_-weighted MRI image of MNP, BV, and MBV solutions and the corresponding T_2_ map; PBS was used as a reference. Dashed red circles indicate the positions of wells not visible in the image. **b**. T_2_ relaxivity of MNP, BV, and MBV calculated from the T_2_ map in **a**. At matched iron concentrations, MBV exhibited slightly higher T_2_ relaxivity than MNP, consistent with the accumulation of MNP on the BV surface.

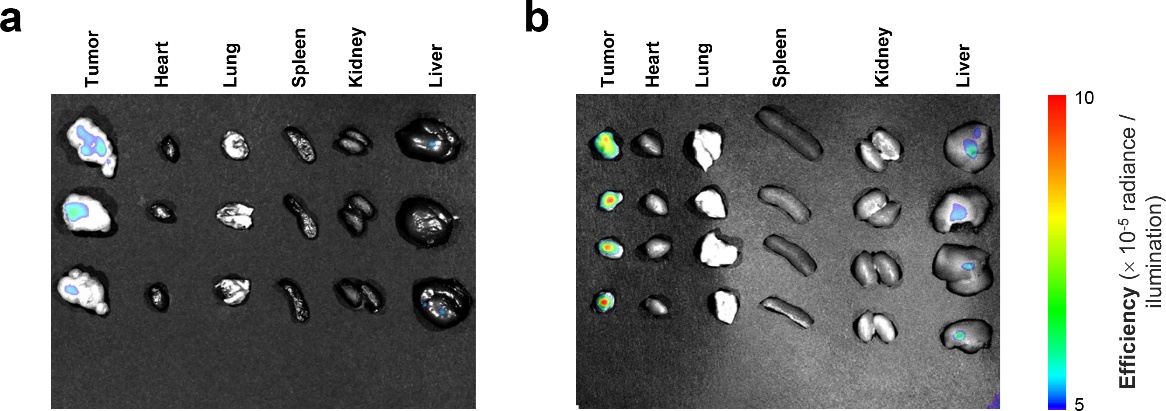

**Fig. S12 | Ex vivo imaging of BV transduction**.

Tumor-bearing mice were treated with PBS (**a**) or MBV–*Pdl1* (**b**) as described in the Methods. 48 h after completion of four treatment, mice were euthanized, and tumors and major organs (heart, lung, spleen, kidney, and liver) were collected and imaged (ex/em = 535/590 nm) using an Ami HT small-animal imaging system.

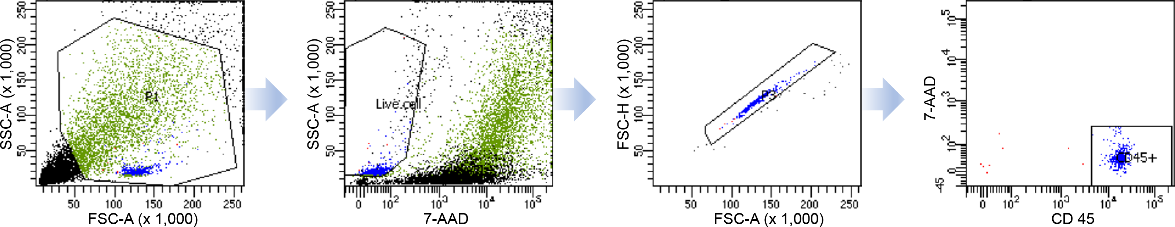

**Fig. S13 | Flow cytometry gating strategy used to isolate tumour-infiltrating immune cells for single-cell RNA sequencing analysis.**

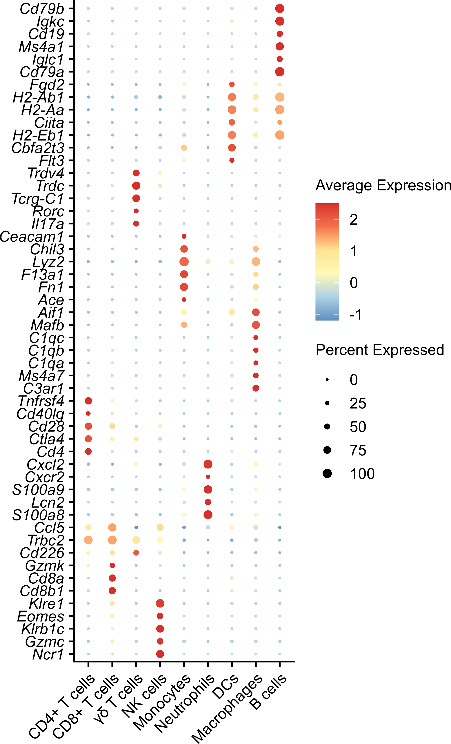

**Fig. S14 | Marker Expression Across Immune Cell Types in single cell RNAseq.**

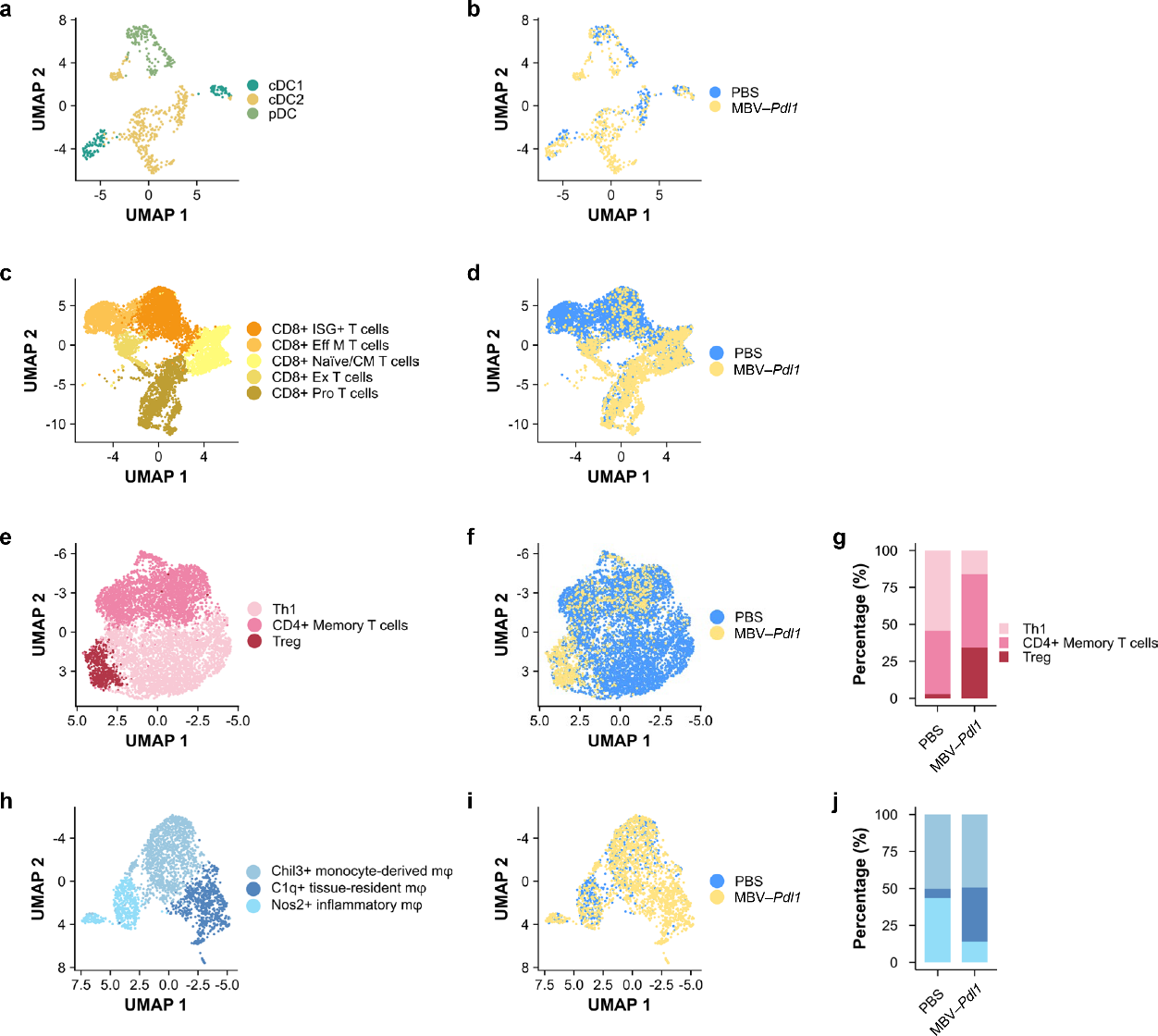

**Fig. S15 | Single cell RNA sequencing of tumor infiltrating immune cells**.

The effects of MBV–*Pdl1* treatment on tumor immune microenvironment were analyzed by single-cell RNA sequencing, in addition to the analyses shown in Fig. 5. **a**. UMAP embedding showing clustering of dendritic cells (DCs). **b**. UMAP showing the distribution of DCs in PBS- and MBV–*Pdl1*-treated tumours. **c**. UMAP embedding showing clustering of CD8^+^ T cells. **o**. UMAP showing the distribution of CD8^+^ T cells in PBS- and MBV–*Pdl1*-treated tumours. **d**. Proportions of CD8^+^ T cell subtypes in PBS- and MBV–*Pdl1*-treated tumours. **e**. UMAP embedding showing clustering of CD4^+^ T cells. **f**. UMAP showing the distribution of CD4^+^ T cells in PBS- and MBV–*Pdl1*-treated tumors. **g**. Proportions of CD4^+^ T cell subtypes in PBS- and MBV–*Pdl1*-treated tumors. **h**. UMAP embedding showing clustering of macrophages. **i**. UMAP showing the distribution of macrophages in PBS- and MBV–*Pdl1*-treated tumors. **j**. Proportions of macrophage subtypes in PBS- and MBV–*Pdl1*-treated tumors.

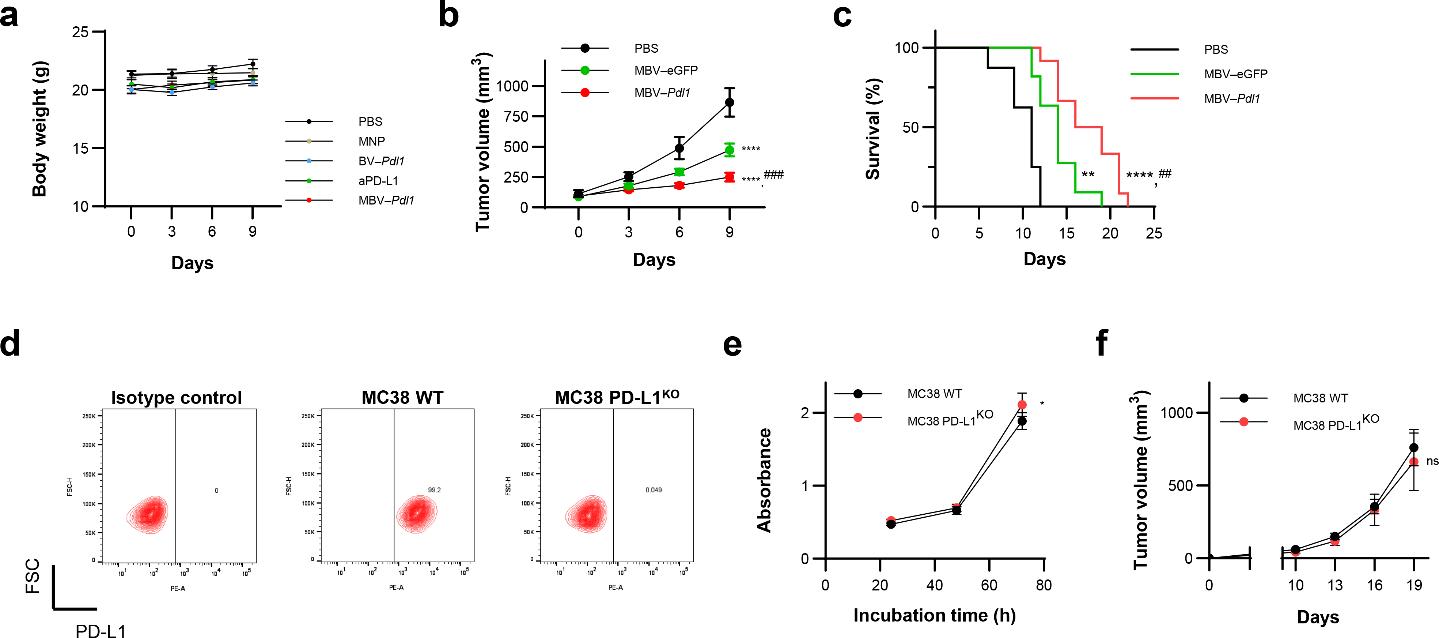

**Fig. S16 | Effects of MBV–*Pdl1* on tumor growth.**

**a**. Body weight monitoring during treatment shown in Fig. 4. To evaluate effects of MBV–eGFP and MBV–*Pdl1* on tumor growth, mice received a subcutaneous injection of 5×10^5^ MC38 cells suspended in 100 μL of PBS into the right flank. Treatments were initiated when tumor volumes reached approximately 50 mm^3^. Mice were randomized to receive PBS, MBV–eGFP, or MBV–*Pdl1*. MBVs were administrated by intratumoral infusion as described in the Methods, except that the MBV concentration was increased fourfold. **b**, **c**. Tumor volume (**b**) and survival curve (**c**). Data are presented as mean ± s.e.m. Sample sizes were n = 8 (PBS), 11 (MBV–eGFP), and 12 (MBV–*Pdl1*). ** and **** denote *P* < 0.01 and *P* < 0.0001 versus PBS, respectively; ## and ### denote *P* < 0.01 and *P* < 0.001 versus MBV–eGFP, respectively. MBV–*Pdl1* treatment resulted in reduced tumor growth and prolonged survival relative to MBV–eGFP, consistent with a contribution from *Pdl1* disruption. To assess the contribution of PD-L1 expression to tumorigenicity, a monoclonal MC38 line with PD-L1 knockout disruption was generated by transfection with a pX330–*Pdl1* plasmid. **d**. Loss of PD-L1 expression confirmed by flow cytometry.**e**. In vitro cell growth measured by CCK-8 assay. Data are presented as mean ± s.d. (n = 4 per group). * denotes *P* < 0.05 versus MC38 WT. **f**. In vivo tumor growth curves. Data are presented as mean ± s.e.m. Sample sizes were n = 8 (MC38 WT) and n = 7 (MC38 PD-L1^KO^). ns denotes no significant difference versus MC38 WT. MC38 PD-L1^KO^ cells exhibited a modest reduction in tumorigenicity (*P* = 0.11, two-tailed paired t test).

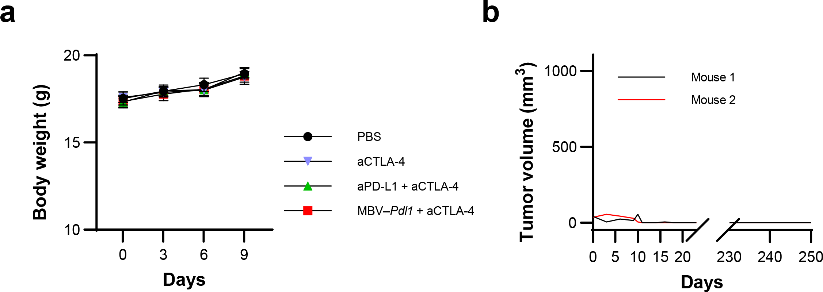

**Fig. S17 | MBV**–***Pdl1* + CTLA-4 therapy.**

**a.** Body weight monitoring during treatment shown in Fig. 6. **b**. Rechallenge of long-term survivors. Tumor growth curves of the two long-term survivors from the MBV–*Pdl1* + anti-CTLA-4 group (Fig. 6), were plotted from initiation of treatment. The x-axis is displayed in two segments: the left segment shows the first 20 days following treatment initiation, and the right segment shows days 230-250, corresponding to the rechallenge period. On day 230, mice were rechallenged by subcutaneous injection of 1 × 10^6^ MC38 cells into the contralateral flank. Tumor growth was monitored for 20 days after implantation. No tumor formation was detected in either mouse.

**Table S1 | List of guide RNAs**

| **gRNA** | **gRNA + NGG** (5’ - 3’) | **Exon number** | **VBC-score** |
| --- | --- | --- | --- |
| VB1 | GTATGGCAGCAACGTCACGATGG | Exon 3 | 0.75 |
| VB2 | GGACCGTGGACACTACAATGAGG | Exon 5 | 0.757 |
| VB3 | GCCAGGGCAAAACCACACAGCGG | Exon 4 | 0.716 |
| VB4 | GGCTCCAAAGGACTTGTACGTGG | Exon 3 | 0.725 |
| VB5 | GCCTGCTGTCACTTGCTACGGGG | Exon 2 | 0.686 |
| VB6 | AGTACACCACTAACGCAAGCAGG | Exon 3 | 0.701 |

**Table S2 | List of primers for T7E1 assay**

| **gRNA** | **Forward** (5’ - 3’) | **Reverse** (5’ - 3’) |
| --- | --- | --- |
| VB1 | AATGAACAACAACCGCCC | CGAACGAATGAACAAACGAG |
| VB2 | CAAGGAAGTTACTGCACTAAGG | GTGTAACTGAAAGCAAGCCC |
| VB3 | GCAGACTAACACTCACTCCC | TCCTATCCAGCCACGAATAC |
| VB4 | AATGAACAACAACCGCCC | CGAACGAATGAACAAACGAG |
| VB5 | CCCATCATACTGACTTCTTTCC | TAGAAGCCAGGTGCAGTAG |
| VB6 | AATGAACAACAACCGCCC | CGAACGAATGAACAAACGAG |

**Table S3 | List of primers for NGS**

### On target

| **gRNA** | **Forward** (5’ - 3’) | **Reverse** (5’ - 3’) |
| --- | --- | --- |
| VB1 | ACACGACGCTCTTCCGATCTTCTGTCTTCTGAGGGCTGGT | GACGTGTGCTCTTCCGATCTTAAGGTCCTCCTCTCCTGCC |
| VB2 | ACACGACGCTCTTCCGATCTACCTTCCATCAGCTTCTCTT | GACGTGTGCTCTTCCGATCTGATGTGGGTCTGTTCTTGTC |
| VB3 | ACACGACGCTCTTCCGATCTAGAGGGGATGCTTCTCAATG | GACGTGTGCTCTTCCGATCTTCGGCCAACTACTGCTAAAT |
| VB4 | ACACGACGCTCTTCCGATCTTCTGTCTTCTGAGGGCTGGT | GACGTGTGCTCTTCCGATCTTAAGGTCCTCCTCTCCTGCC |
| VB5 | ACACGACGCTCTTCCGATCTAGACCTCTCTGTGTTTCCCG | GACGTGTGCTCTTCCGATCTCGCTTTATGTGTGCCTAGCA |
| VB6 | ACACGACGCTCTTCCGATCTTATGGCAGCAACGTCACGAT | GACGTGTGCTCTTCCGATCTTGCAGCTTGACGTCTGTGAT |

### Off target

| **gRNA** | **Forward** (5’ - 3’) | **Reverse** (5’ - 3’) |
| --- | --- | --- |
| VB1-1 | ACACGACGCTCTTCCGATCTTCCCGAGCCTTACGTACAGA | GACGTGTGCTCTTCCGATCTGCACGGGTCTCTCCATTCTC |
| VB1-2 | ACACGACGCTCTTCCGATCTTGGCTAACATTTAAAGGCAC | GACGTGTGCTCTTCCGATCTTGGGAAGAAAGTCCAACTCA |
| VB1-3 | ACACGACGCTCTTCCGATCTGTGAGCCACAAGGAAGTATG | GACGTGTGCTCTTCCGATCTCCATCCAACCCACAGAAAAT |
| VB1-4 | ACACGACGCTCTTCCGATCTAGTCATCCTCCACACCCAGC | GACGTGTGCTCTTCCGATCTGCTGCCTCCCAGGTAACCTT |
| VB2-1 | ACACGACGCTCTTCCGATCTTCCACATTGGAGCATCAGCA | GACGTGTGCTCTTCCGATCTAGTGTGTTTTGAAAAGAAGGGAGA |
| VB2-2 | ACACGACGCTCTTCCGATCTTCTTTGGGCCTCTTCAGCAG | GACGTGTGCTCTTCCGATCTCGTCACCCATAATCCCCTGG |
| VB2-3 | ACACGACGCTCTTCCGATCTATATGGGCAACAATTGGACC | GACGTGTGCTCTTCCGATCTCAGCATCACTGTTCAAGACA |
| VB2-4 | ACACGACGCTCTTCCGATCTAGATTTACTACCCCAGAGCC | GACGTGTGCTCTTCCGATCTATGTACAACCATCTGGGACA |
| VB3-1 | ACACGACGCTCTTCCGATCTACCATGGGTTATGATGGCCA | GACGTGTGCTCTTCCGATCTGGAGAACCCATTACGACGGT |
| VB3-2 | ACACGACGCTCTTCCGATCTCTCAGAAGACCACGAGGCAG | GACGTGTGCTCTTCCGATCTGCATCCCTGGGAATATGCCA |
| VB3-3 | ACACGACGCTCTTCCGATCTGCAGTTCACTTCATAGTACATGG | GACGTGTGCTCTTCCGATCTAATGCACGTGACAGAGATGT |
| VB3-4 | ACACGACGCTCTTCCGATCTGCTGGCCTCCATCTCCTGAT | GACGTGTGCTCTTCCGATCTTCTTACAGAGTGGGACCAGCA |
| VB4-1 | ACACGACGCTCTTCCGATCTGCTAGCCATCCCACGGAAAA | GACGTGTGCTCTTCCGATCTGGACGTTCTCTGTCTGCCCT |
| VB4-2 | ACACGACGCTCTTCCGATCTGGGGAATCCTGACCCTGCAA | GACGTGTGCTCTTCCGATCTTCCGCTAGTGGGAATTCAGC |
| VB4-3 | ACACGACGCTCTTCCGATCTGCTTCGAGTGACTCCAGCTT | GACGTGTGCTCTTCCGATCTGGCTAAGGGCAATGGGACAT |
| VB4-4 | ACACGACGCTCTTCCGATCTACAAGCTGCTGAAAACCTGG | GACGTGTGCTCTTCCGATCTCCTCTGAAGAAGGCTCACGA |
| VB5-1 | ACACGACGCTCTTCCGATCTCCTGAACCCACGCCCCATAA | GACGTGTGCTCTTCCGATCTTGTCACCCTTGCCTCTCTTGG |
| VB5-2 | ACACGACGCTCTTCCGATCTCTGAGATCGTCTGCAGACCC | GACGTGTGCTCTTCCGATCTCAGCCCAGGGACAGTATTGG |
| VB5-3 | ACACGACGCTCTTCCGATCTGCAGTTTGTCTGTGTCCTGG | GACGTGTGCTCTTCCGATCTTGACAGGGGTCCTTTTCTCC |
| VB5-4 | ACACGACGCTCTTCCGATCTAACTGTGTGCTGGTGTGACA | GACGTGTGCTCTTCCGATCTTTTTCCTGTGACGGACGTGC |
| VB6-1 | ACACGACGCTCTTCCGATCTCACTTCTCTGCTGGCTGGAA | GACGTGTGCTCTTCCGATCTTGTCAAAGCCATGCTCTCGT |
| VB6-2 | ACACGACGCTCTTCCGATCTACGAGAGGCACATGTTGAGA | GACGTGTGCTCTTCCGATCTACAATGATGCTGCCTTCCTCT |
| VB6-3 | ACACGACGCTCTTCCGATCTGAACTGGCTTGCGTAGACCT | GACGTGTGCTCTTCCGATCTTACCCAGAGCTAGCCTCCTG |
| VB6-4 | ACACGACGCTCTTCCGATCTTCTGCAACCAAACTGGGCTT | GACGTGTGCTCTTCCGATCTGCTAGACTTTGGCAGGAACCT |

**Table S4 | List of primers for RT-qPCR**

| **Gene** | **Forward** (5’ - 3’) | **Reverse** (5’ - 3’) |
| --- | --- | --- |
| GAPDH | GCATCTTCTTGTGCAGTGCC | ACTGTGCCGTTGAATTTGCC |
| CD86 | ACGGACTTGAACAACCAGACT | CGTCTCCACGGAAACAGCAT |
| CD40 | CTGCTGGTCATTCCTGTCGT | GTTCCAGGGTTCAGACCAGG |
| IL-12 | TAGAGGTGGACTGGACTCCC | GTGAGTGGCTCAGAGTCTCG |
| TNF-α | CAGTTCTATGGCCCAGACCC | TAGCAAATCGGCTGACGGTG |
| IFN-α1 | AGGACTTTGGATTCCCGCAG | TCATTGAGCTGCTGGTGGAG |
| IFN-β1 | CAACCTCACCTACAGGGCG | CTGTCTGCTGGTGGAGTTCAT |
| GranB | AGGACTTTGTGCTGACTGCT | TCACATTGACATTGCGCCTG |
| IFN-γ | CGGCACAGTCATTGAAAGCC | TGCATCCTTTTTCGCCTTGC |
| IL-2 | GAAACTCCCCAGGATGCTCA | AAAGTCCACCACAGTTGCTGA |
| IL-10 | GGTTGCCAAGCCTTATCGGA | ACACCTTGGTCTTGGAGCTTA |

**Table S5 | List of antibodies.**

| **Name** | **Vendor** | **Catalog no.** |
| --- | --- | --- |
| InVivoMAb anti-mouse PD-L1 (clone 10F.9G2™) | Bio X Cell | BE0101 |
| InVivoMAb anti-mouse CTLA-4 (clone 9H10) | Bio X Cell | BE0131 |
| Alexa Fluor® 594-conjugated rat anti-mouse CD3 | Biolegend | 100240 |
| APC-conjugated rat anti-mouse CD45 | Biolegend | 103112 |
| PE/Cyanine-conjugated rat anti-mouse CD44 | Biolegend | 163607 |
| PE-conjugated rat anti-mouse PD-L1 | Biolegend | 124308 |
| Alexa Fluor® 594-conjugated hamster anti-mouse CD11c | Biolegend | 117346 |
| Rat anti-mouse CD86 | Abcam | ab119857 |
| Rabbit anti-mouse CD8a | Abcam | ab217344 |
| Alexa Fluor® 555-conjugated goat anti rat antibody | Abcam | ab150158 |
| Alexa Fluor® 488-conjugated goat anti rabbit antibody | Abcam | ab150077 |
| Alexa Fluor® 647-conjugated rat anti-mouse ER-TR7 | Santa Cruz Biotechnology | Sc-73355 |
